# Genes and Pathways Comprising the Human and Mouse ORFeomes Display Distinct Codon Bias Signatures that Can Regulate Protein Levels

**DOI:** 10.1101/2025.02.03.636209

**Authors:** Evan T. Davis, Rahul Raman, Shane R. Byrne, Farzan Ghanegolmohammadi, Chetna Mathur, Ulrike Begley, Peter C. Dedon, Thomas J. Begley

**Author notes:** Codomax, Massachusetts Biomedical Initiatives (MBI), 17 Briden St, STE 220, Worcester, MA 01605. Astellas Institute for Regenerative Medicine, 9 Technology Dr, Westborough, MA 01581, USA. Corresponding Authors: Peter C. Dedon, and Thomas J. Begley. Declarations: SRB, PCD, and TJB hold equity in Codomax.

## Abstract

Arginine, glutamic acid and selenocysteine based codon bias has been shown to regulate the translation of specific mRNAs for proteins that participate in stress responses, cell cycle and transcriptional regulation. Defining codon-bias in gene networks has the potential to identify other pathways under translational control. Here we have used computational methods to analyze the ORFeome of all unique human (19,711) and mouse (22,138) open-reading frames (ORFs) to characterize codon-usage and codon-bias in genes and biological processes. We show that ORFeome-wide clustering of gene-specific codon frequency data can be used to identify ontology-enriched biological processes and gene networks, with developmental and immunological programs well represented for both humans and mice. We developed codon over-use ontology mapping and hierarchical clustering to identify multi-codon bias signatures in human and mouse genes linked to signaling, development, mitochondria and metabolism, among others. The most distinct multi-codon bias signatures were identified in human genes linked to skin development and RNA metabolism, and in mouse genes linked to olfactory transduction and ribosome, highlighting species-specific pathways potentially regulated by translation. Extreme codon bias was identified in genes that included transcription factors and histone variants. We show that re-engineering extreme usage of C- or U-ending codons for aspartic acid, asparagine, histidine and tyrosine in the transcription factors *CEBPB* and *MIER1,* respectively, significantly regulates protein levels. Our study highlights that multi-codon bias signatures can be linked to specific biological pathways and that extreme codon bias with regulatory potential exists in transcription factors for immune response and development.

**Graphical Abstract:** 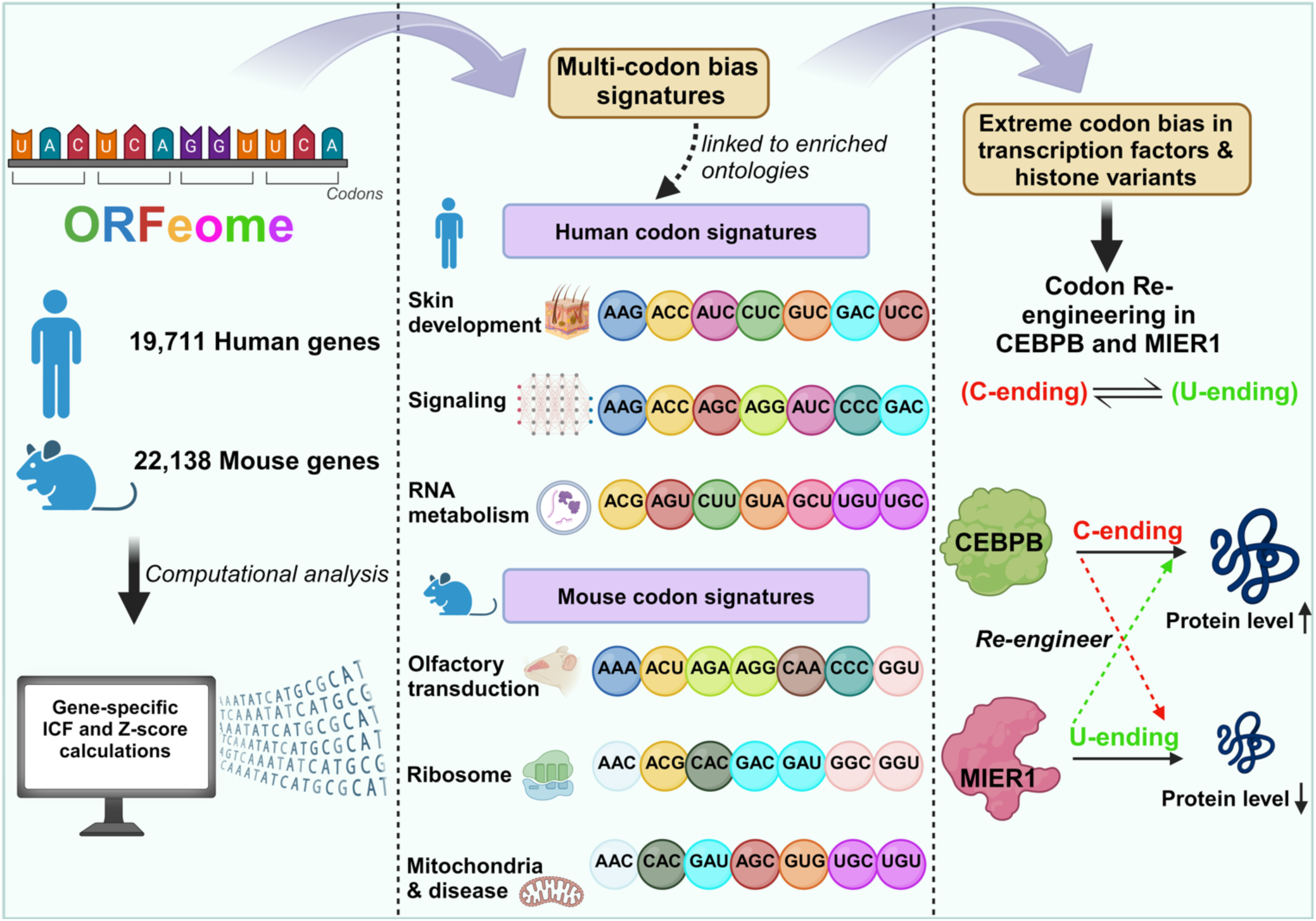

## Introduction

The cytoplasmic translational machinery can use 1 to 6 synonymous codons to decode each of 21 amino acids. Genome sequencing projects have provided significant data to characterize codon usage using metrics that include Isoacceptor Codon Frequencies (ICF), Total Codon Frequency (TCF), Relative Synonymous Codon Usage (RSCU), Codon Adaptation Index (CAI), tRNA Adaptation Index (tAI) and Supply Demand Adaptation (SDA), among many other measures.^1–5^ Codon usage metrics were broadly developed to answer questions related to evolution or protein synthesis, with some also utilizing tRNA gene or expression levels to inform on translation.^1^ Most codon usage metrics detail or can be processed to describe gene-based codon usage bias, which in some cases describe a preference for specific synonymous codons for an amino acid. The functional genomic rules governing codon bias and genome-wide trends are still being determined and are an important area of research.^3,6–9^ Synonymous codons have inherent translational regulatory features due to their sequence variations at the third position, also known as the mRNA wobble base. Studies using reporter or synthetic genes have demonstrated that swapping synonymous codons can alter protein levels significantly (250-fold for GFP, other)^6,10^, but maintaining a balance between codon usage and tRNA abundance is an important parameter for maintaining efficient translation.

Classically, the efficiency of codon decoding was attributed to the pool of available cognate tRNAs, and codon usage was considered a static metric. Positions 34 and 37 of the tRNA anticodon are hot spots for RNA modifications that can regulate codon-anticodon interactions. Multiple studies have recently demonstrated that the tRNA epitranscriptome can be dynamically re-programmed in response to changing cellular physiology and stress, and shown that codon usage preference and decoding can be modulated during cellular responses.^11–21^ tRNA modifications whose decoding potential matches the codon biased mRNAs are key regulators of translation, with the translationally regulated mRNAs termed modification tunable transcripts (MoTTS).^22^ In yeast, pathways enriched with MoTTs have been identified using ORFeome wide gene-specific codon bias measures and comprise mRNAs encoding proteins participating in DNA damage and stress response, protein synthesis, energy and metabolism.^15,17,23^

Here we have used ORFeome wide analysis and comparison of codon usage and bias data between humans and mice to identify similar and distinct pathways housing potential MoTTs. We have shown that there are global similarities in codon usage between humans and mice and some overlapping biological processes that have distinct codon-bias signatures. Each species has exploited codon bias in different pathways though, with humans using multi-codon bias signatures in genes linked to skin development and mice using it for olfactory transduction genes linked to smell, among others. We have also demonstrated that extreme codon bias can be identified in human and mice genes, with mRNAs for some transcription factors (i.e., *CEBPB* and *MIER1*) and histone proteins totally or mostly committed to specific synonymous codons. We used re-engineering of the extremely codon biased *CEBPB* and *MIER1* to demonstrate that exchanging C-ending for -U ending synonymous codons for asparagine, aspartic acid, histidine and tyrosine, and vice versa, can dramatically regulate protein levels and highlight extreme codon usage as potential regulatory mechanism for transcription factors.

## Materials and Methods

### Codon Counting, Frequency and Z-score calculations

Complete open reading frames (ORFs) for all human coding sequences was downloaded from NCBI (GRCh38) at https://ftp.ncbi.nlm.nih.gov/genomes/refseq/vertebrate_mammalian/Homo_sapiens/all_assembly_versions/GCF_000001405.39_GRCh38.p13/.

Mouse coding sequences were downloaded from NCBI (GRCm39) at https://ftp.ncbi.nlm.nih.gov/genomes/refseq/vertebrate_mammalian/Mus_musculus/all_assembly_versions/GCF_000001635.27_GRCm39/.

Gene sequences were analyzed using our gene specific codon counting (GSCU) algorithm,^15,17,24^ to obtain codon counts, ICF and Z-scores. Briefly, ICF inform on the use of a synonymous codon for a specific amino acid, with the number of synonymous codons ranging from 2 to 6 for each amino acid. ICF was chosen as it has proven to predict protein levels during stress responses, is a driver of codon biased translational regulation observed in many species and can normalize for amino acid bias.^8,13,15,17,23–30^ Z-scores detail whether a gene is over- or under-using a synonymous codon for a specific amino acid, relative to genome averages. Corresponding codon data for all human and mouse genes was compiled and analyzed using Python based methods, as described below and available on Github.

### Scripts to Characterize Codon Bias

All analysis was performed using Python 3.8.5, Jupyter 1.0.0, and Pandas 1.2.3. ICF and Z-score plots were generated using matplotlib 3.3.4. All code can be found and accessed in the below folder. https://www.dropbox.com/scl/fo/f79mi1damm7d01y4rhvoo/AIA-bsagh-VLYEF18_Jh5xo?rlkey=9hktrl4vocj1pv3duqcu3ka7b&st=qxb28bhq&dl=0

#### Number of codons enriched in each gene data matrix

A gene-specific multi-codon Z-score compiler function analyzed all genes in a species, generated a list of genes over- or under-using a specific codon at the specified Z-score threshold, iterated this process for all codons, and then was collated to generate a dictionary of genes over-using multiple codons. The dictionary was then used to identify the number of genes over- or under-using N codons (N = 2 to 62) at a specified Z-score threshold. Gene lists were specified from the dictionary and then analyzed using STRING to perform gene ontology analysis.^31^ The gene-specific multi-codon Z-score compiler function was also used to generate a dictionary detailing the number of genes that over- or under-use each specific codon (N = 62) at discrete Z-score thresholds (Z=> 1.0, 1.5, 2.0, 2.5, 3.0, 3.5, 4.0, 4.5, 5.0 and Z <= -1.0, -1.5, -2.0, -2.5, -3.0, -3.5, -4.0, -4.5 and -5.0). Codon specific data tables were developed from the dictionary detailing the number of genes meeting the Z-score threshold for all codons, with the corresponding data matrix used to generate heatmap in Morpheus.^32^

#### Codon over-use ontology mapping

Lists describing genes that over-use a specific codon (Z => 2) were assessed for gene ontology enrichment (biological and KEGG functions using Selenium 3.141.0 to access the STRING database). If a term description was observed in more than one gene list, the associated codon, term description and FDR value were used to construct a data matrix. Once all codons and term descriptors were retrieved, the -*log*_10_(FDR) was calculated for each value and the resulting table was used to construct a heatmap in Morpheus with hierarchical clustering on rows and columns.

#### Gene-specific summed Z-score (GSZ-score) calculation

The absolute value of each codon-specific Z-score for each gene was used to calculate the GSZ-score. Genes were then sorted to generate gene lists for the top 50, 1% and 2.5% scoring genes and then analyzed in the STRING database for gene ontology enrichment. In addition, the ICF for each codon for the top 50 scoring genes were analyzed using hierarchical clustering in Morpheus.^32^

#### Cell studies and gene engineering

HepG2 cells were seeded in 6-well plates at 5 x 10^5^ cells/well and transfected with TransfeX™ Transfection Reagent (ATCC, Manassas, VA) with pCMV 3xFLAG vector (Agilent, Santa Clara, CA) expressing either engineered *CEPBP* or *MIER1* constructs. Genes were synthesized and cloned by Genescript (Piscataway NJ). Briefly, 2.5 μg of plasmid DNA and 5 μl of Transfection Reagent, were diluted in 250 μl of Opti-MEM™ Reduced Serum Medium (Gibco, Thermo Fisher Scientific, Waltham, MA) according to manufacturer’s protocol. For CEBPB studies, 24 H after transfection cells were left untreated or treated with 0.12 mM [LD_20_] NaAsO_2_ (Sigma-Aldrich, St. Louis, MO) for 2 hours in complete growth media. Cells were then harvested after 48 hours of transfection and lysed in RIPA buffer (50mM Tris-HCl pH 7.4. 150mM NaCl, 1% Triton-X 100, 1% Sodium deoxycholate, 0.1% SDS, 1mM EDTA with Protease Inhibitors) at 4 °C for 30 min, after which the lysates were cleared of cell debris by using centrifugation at 2000× *g* for 5 min. The protein concentrations of the samples were quantitated using Bradford Protein Assay (Bio-Rad, Hercules, CA). Protein lysates were analyzed using WES Simple Western™ instrument (ProteinSimple^®^, Bio-Techne, Minneapolis, MN). Samples were mixed with 1x fluorescent master mix (EZ standard pack I; ProteinSimple^®^) according to protocol and 2.88 µg total protein in a 3 µl volume was loaded into each well. Monoclonal ANTI-FLAG^®^ M2 antibody produced in mouse (Sigma-Aldrich, St. Louis, MO) was used at a 1:1000 dilution in Milk-Free Ab diluent (Bio-Techne), whereas the loading control Anti-Neomycin Phosphotransferase II Antibody produced in rabbit (Sigma-Aldrich, St. Louis, MO) was used at a 1:10 dilution in Milk-Free Ab diluent (Bio-Techne). The secondary antibodies (anti-mouse and anti-rabbit HRP) and enhanced chemiluminescence (ECL) reagents were used according to the kit’s instructions (ProteinSimple^®^, Bio-Techne). Either the 13 or 25 capillary cartridges (12–230 kDa separation module, ProteinSimple^®^, Bio-Techne) were used for protein analysis. WES Simple Western™ data were analyzed using Compass for Simple Western software.

### Calculation of free energy RNA structures for wild-type and codon engineered constructs

RNA structure calculations detailing minimum free energy structure (MFE) of the wild-type and codon engineered constructs of *CEPBP* and *MIER1* were calculated using three different tools; RNAfold (http://rna.tbi.univie.ac.at/cgi-bin/RNAWebSuite/RNAfold.cgi), UNAfold (http://www.unafold.org/mfold/applications/rna-folding-form.php) and Sfold (https://sfold.wadsworth.org/cgi-bin/srna.pl).^33–36^ In the case of RNAfold, the following default parameters were used: no folding constraints specified, avoid isolated base pairs, dangling energies on both sides of the helix, RNA parameters from the 2004 Turner model, rescaling energy parameters to 37°C and using 1M salt concentration. In the case of UNAfold, the default parameters were the same temperature and salt concentration as that used by RNAfold and maximum of 50 computed folded structures with maximum interior bulge loop size and asymmetry set to 30. The same default temperature of 37°C and salt concentration of 1 M was used in Sfold server.

## Results

### Each codon displays distinct gene-specific patterns of usage in the human and mouse ORFeome

We generated ICF and Z-scores **(Fig. 1A-B)** for 62 codons in each of 19,711 human and 22,138 mouse genes comprising their respective ORFeomes (**Supplemental Fig. S1, Supplemental Tables S1)**. ICF values describe if a synonymous codon is preferred in a gene sequence, and it is a measure that normalizes for amnio acid bias. For each codon we binned the number of genes that had a codon-specific ICF range from 0 to 1, at 0.1 intervals, to identify codons with distinct usage characteristics **(Fig. 1C, left)**. Z-scores detail how many standard deviations away from the genomic mean a codon frequency is in a specific gene, and we have labeled Z=> 2 as over-use and Z=< -2 as under-use. Z-score histograms were generated to identify the number of genes that over- or under-use a codon, as well as the distribution **(Fig. 1C, right).** Genes over-using a codon (Z >= 2 or 4) were identified in humans and mice for 62 codons **(Supplemental Tables S2)**. 15,300 genes in humans and 15,284 genes in mice overuse 1 - 2 codons with a Z => 2. 3,562 human genes and 3,258 mouse genes over-use 1 - 2 codons when we increased Z => 4. We identified 389 human genes (N = 13 or more codons) and 367 mouse genes (N = 12 or more codons) that over-use N-codons at a Z =>2, and the corresponding gene lists were analyzed for gene ontology enrichments **(Supplemental Table S3)**. Both organism-specific gene lists over-using N-codons (13 for humans and 12 for mice) showed enrichment of mitochondrial related genes, which should be expected as the mitochondria is an A/T rich genome relative to the nuclear genome. The mouse ontologies of beta defensin and Parkinson’s disease were also identified.

**Figure 1.**
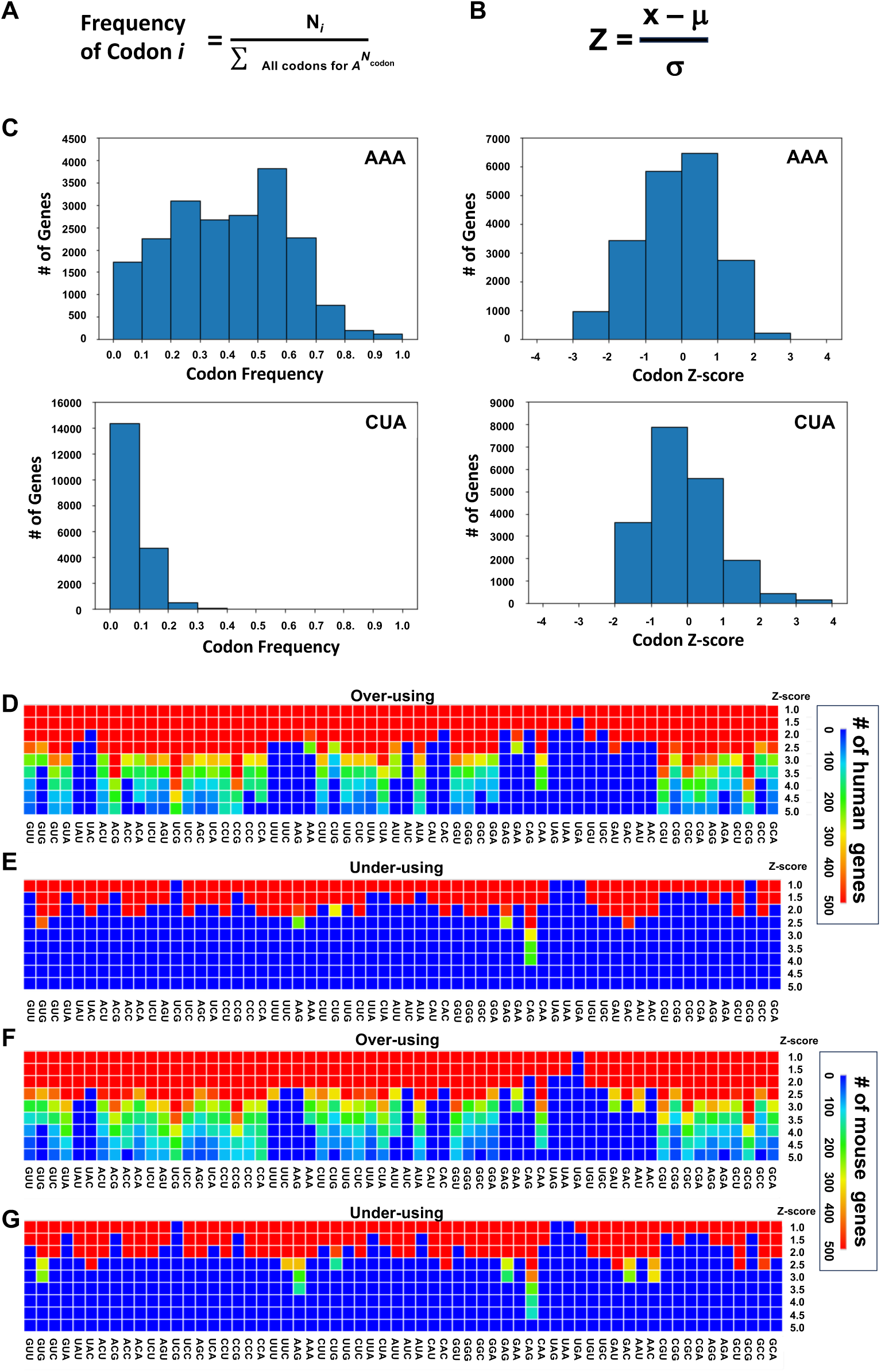
ORFeome Wide Codon Metrics for Human Genes. **A.** Gene-specific ICF formula, where N*_i_* is the number of times codon *i* appears in the mRNA sequence. ∑all codons for 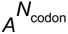 is the sum of all occurrences of codons that encode that amino acid A. **B.** Gene-specific codon Z-score formula uses frequency (x), average frequency of the genome (μ), and standard deviation of the genome average (σ). **C.** Representative plot for human genes at each codon frequency (left) or Z-score (right) for AAA (top) and CTA (bottom). The number of genes over-using or under-using a codon at different Z-score thresholds in humans **(D-E)** and mice **(F-G)**.

### Human and mouse genes have overlapping and species-specific patterns of codon bias

Heatmaps were used to visualize the number of genes that over- or under-use each codon at different Z-score thresholds for both the human and mouse data sets **(Fig. 1D-G, Supplemental Tables S4A-D)**. As Z-score thresholds were increased fewer human and mouse genes over-use a codon **(Fig. 1D & F)**. We identified 200+ genes (green color) in both species over-using the UCG (Ser) codon at a Z => 5. Using a Z-score => 3.0 we identified 8 codons in humans and in 8 mice that were over-used in => 200 genes **(Fig. 1D & F)** and they were GCG (Ala), CCA (Pro), CCG (Pro), UCG (Ser), CGA (Arg), CGU (Arg), CUA (Leu) and ACG (Thr). There were fewer under-used codons in the analysis of human and mouse genes, with neither species having any genes with Z =< -5. **(Fig. 1E & G)**. Interestingly for both humans and mice, the codons CAG (Gln), AAG (Lys) and GAG (Glu) were distinctly under-used in some genes (Z < = -2.5, gene counts => 200). In addition, mouse genes more frequently under-use the GUG (Val), GAC (Asp) and AAC (Asn) codons relative to humans. Our findings highlight that some codons bias is generally conserved between humans and mice, with some species-specific differences in the extent (*i.e.*, number of genes meeting Z-score thresholds) of codon bias.

### Gene-specific ICF highlight a stratified ORFeome enriched in distinct biological networks

Gene-specific ICF data for each of 59 codons was used to stratify the 19,711 human **(Fig. 2, Supplemental Table S5A)** and 22,138 mouse genes **(Fig. 3, Supplemental Table S5B)**. All species specific ICF data was hierarchically clustered and visualized, with clear patterns of codon usage observed in large groups of genes for both humans and mice **(Fig. 2 & 3)**. The patterning in the human verse mouse map is different **(Fig. 2 & Fig. 3)**, with humans having 5 (+2 sub-groups) and mice having 8 distinct clusters. For humans, a distinct codon pattern highlights the Group A cluster with ICF values between ∼0.6 and 0.8 for U/A-ending codons **(Fig. 2)** for UGU (Cys), CAU (His), AAU (Asn), GAA (Glu), AAA (Lys), UUU (Phe) and UAU (Tyr). Human group A is the extreme U/A-ending cluster. The human Group B cluster had decreased ICF values for codons described for group A (∼0.4 to 0.6), but also had increased ICF values (∼0.6 to 0.8) for many C/G-ending codons **(Fig. 2)**. Human group B is the intermediary C/G/U/A-ending group. The human Group C cluster has high ICF-values for C-ending codons and intermediate ICF-values for U-ending codons **(Fig. 2)**, with this cluster being a more extreme variation of group B. Human Group D has the highest ICF values for many C- and G-ending codons and is the extreme C/G-ending cluster **(Fig. 2)**. Human group E is a small group of genes with ICF values in ∼0.5 to 0.8 ranges for many U/A-ending codons and very low values ∼0.0 to 0.2 for C/G-ending codons **(Fig. 2)**. Group E genes are linked to the mitochondria. Some specific sub-groups for B and D (B1 & D1) were also identified as distinct codon users. Sub-group B1 was enriched for processes linked to the detection of chemical stimulus and immune system components. Subgroup D1 had the highest CGC (Arg) ICF values (∼0.4 to 0.8) in the genome and was enriched in development process linked to tissues, cells and the nervous system.

**Figure 2.**
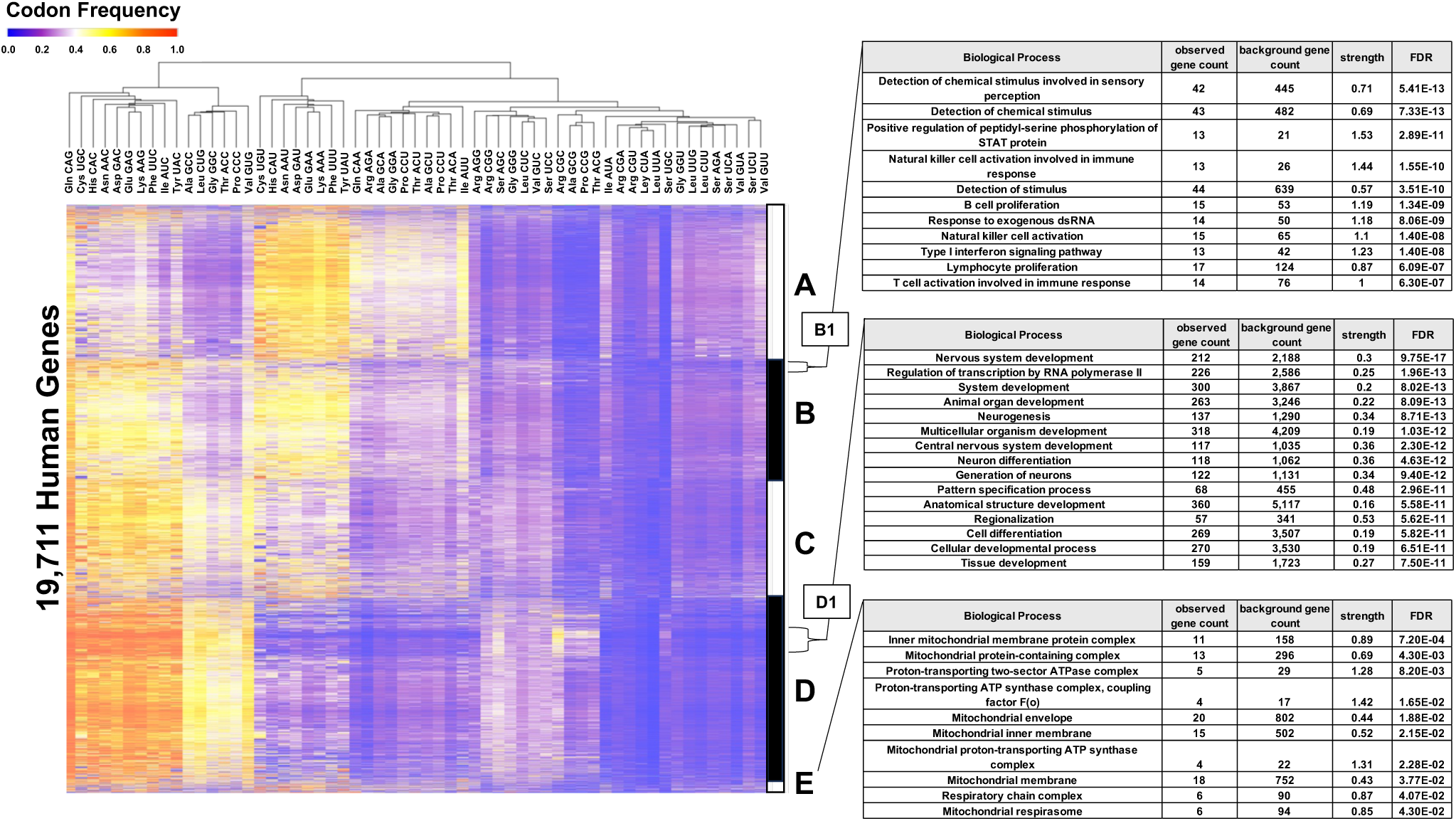
Gene-Specific Codon Frequency Maps for the Human ORFeome. ICF data for 19,711 human genes was clustered into five groups (A-E). Sub-patterns of codon usage were analyzed for enriched gene ontology.

**Figure 3.**
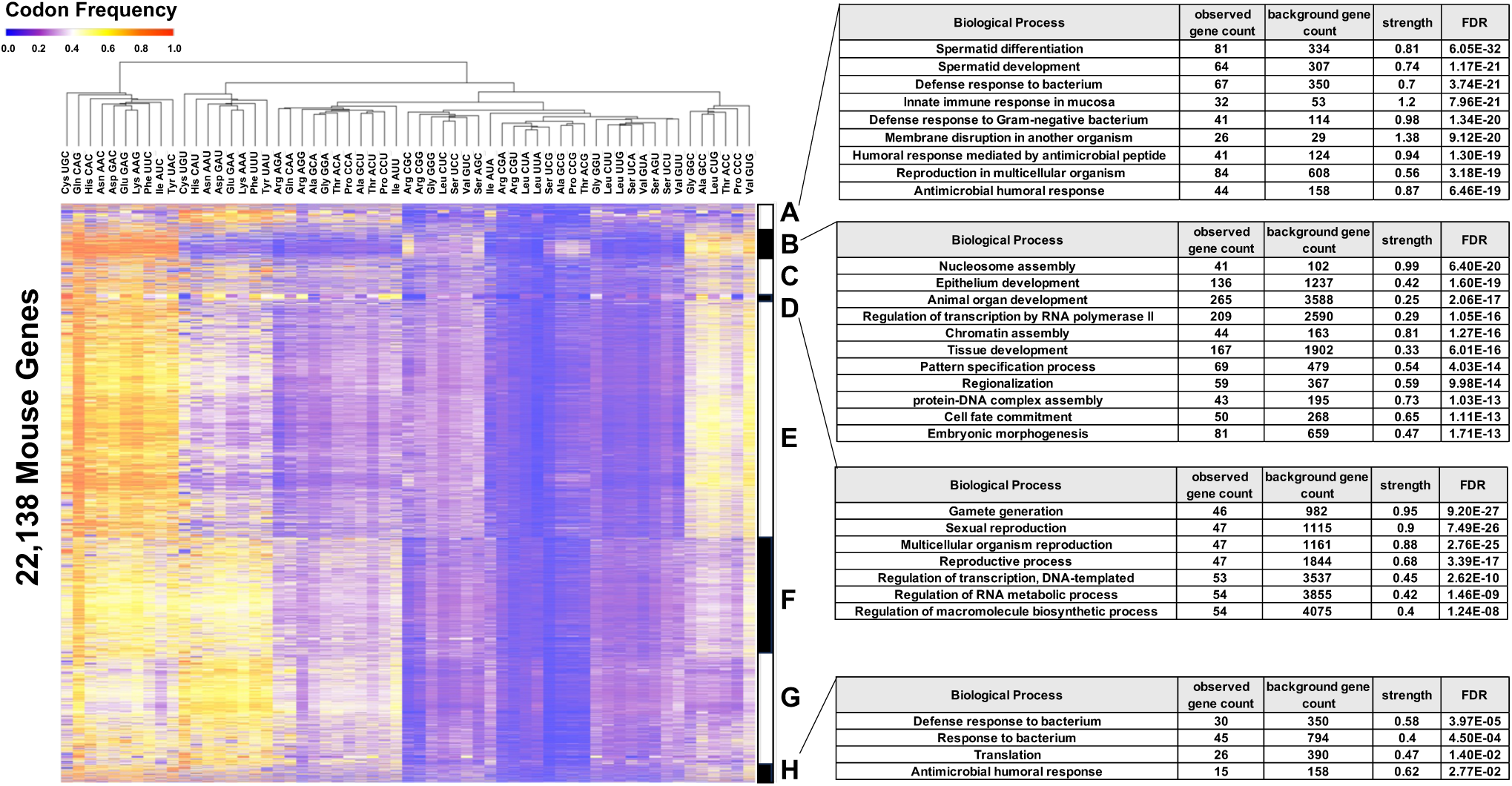
Gene-Specific Codon Frequency Maps for the Mouse ORFeome. ICF data for 22,138 unique mouse genes. Eight general clusters were identified (A-H) representing different patterns of gene-specific codon usage. Enriched gene-ontologies for groups A, B, D and H were identified.

We also analyzed the 22,138 mouse **(Fig. 3)** genes to show that ICF values could be used to stratify into codon-defined patterns. Mouse gene-specific ICF data stratified more groups than the human data (8 groups of A - H vs 5 groups of A - E), but there were some similarities between the two species. Both species had genes that use the CAG (Gln) codon at very high ICF levels, relative to its synonymous partner CAA (Gln). In addition, variations of high and low ICF values for C/G-ending and U/A-ending codons are prime drivers for stratification in both species. Specific to mice, high ICF values (∼0.6 to 0.9) for some U- and A-ending codons highlight a mouse group A representing an extreme U/A-ending group **(Fig. 3)**. Mouse group A genes are linked to spermatid, reproduction and innate immune responses. Mouse group B genes have high ICF values (∼0.7 to 1.0) for specific C- or G-ending codons **(Fig. 3)**. The biological processes of chromatin assembly, development (epithelium, animal organ, and tissue), and transcriptional regulation are linked to mouse group B, with it being an extreme C/G group. There were ontological similarities related to development between mouse group B and human group D1. Mouse group C is a less extreme version of group B. Mouse group D has the most distinct pattern in the heat map and has very low ICF values for many codons, notably Cys UGU (< 0.2) which contrasted by an extremely high (> 0.7) ICF value for Cys UGC. Mouse group D represents genes linked to biological processes related to sexual reproduction and RNA synthesis **(Fig. 3)**. Other ICF defining groups were also identified in mice **(Fig. 3, groups E-H)**. Notably ICF values of ∼0.3 to 0.6 for GGC (Gly), GCC (Ala), CUG (Leu), ACC (Thr), CCC (Pro) and GUG (Val) define mouse group H, which is enriched for biological process linked to innate immunity and translation.

### Codon over-usage patterns are linked to distinct biological functions

We and others have previously reported that codon over-use can be regulatory, and that specific codon biases in the mRNA of functionally related proteins supports a mechanism of translational regulation where key tRNA modifications regulate protein synthesis.^8,13–15,23,24,26,28^ Our goal here was to determine if any ontologies were comprised of genes that over-use a specific codon, and then determine if these ontologies could be linked to multiple codons and have a codon bias signature. We began codon over-use ontology mapping by generating 59 lists of human or mouse genes over-using (Z => 2) each codon **(Fig. 4A)**. We analyzed each of 59 gene lists for enriched ontologies and detailed biological processes enriched in only 1 gene list (*i.e.*, linked to only 1 codon) **(Supplemental Tables S6A-B)**. We also identified ontologies that were identified in => 2 lists of genes over-using a specific codon **(Supplemental Tables 7A-B)**. The False Discovery Rate (-*log*_10_ of FDR) data for each codon-linked ontology was recorded in a matrix and clustered and visualized as heatmaps for humans **(Fig. 4B)** and mice **(Fig. 5)** to highlight gene ontologies linked to patterns of multi-codon bias. In the heat map specific to humans **(Fig. 4B)**, ontologies represented by the general descriptors signaling & development, skin differentiation, ion homeostasis, antimicrobial defense & immune response, mitochondria & disease, nucleoside and transport, mRNA, metabolism, translation & targeting, detection of chemical stimulus, and cell cycle are linked to specific patterns of multi-codon bias, with the exact ontologies numerous and shown in Supplemental Figure S2. In humans, gene ontologies whose gene list are enriched with seven or more codons are highlighted by regulation of signaling receptor activity, keratinization and skin development **(Supplemental Table 8A)**. Ontologies that included keratinization, skin development, and epithelial cell differentiation, among others, are grouped under the heading skin differentiation **(Fig. 4B)** and have their corresponding genes over-using many codons, most notably ACC (Thr), CTG (Leu), GAC (Asp) and GCG (Ala), among others. The metabolism **(Fig. 4B)** cluster is dominated by the corresponding genes that over-use AUU (Ile), ACU (Thr) and CUU (Leu) codons, among others, and includes many ontologies similar to biosynthetic processes, RNA metabolic processes, macromolecular metabolic processes and nitrogen compound metabolic processes. The detection of chemical stimulus cluster is linked to smell and includes the ontologies olfactory transduction, sensory perception of smell, and detection of chemical stimulus, with the corresponding genes primarily over using AGG (Lys) and UCG (Ser) codons. We also parsed the input list to identify only those ontologies linked to gene lists that overuse at least 5 distinct codons (-*log*_10_ of FDR values > 2), clustered this restricted list and visualized as a heatmap **(Fig. 4C)**. The two most codon distinct human ontologies are (1) regulation of signaling receptor activity and (2) keratinization, as they are both are linked to 6 codon-defined gene lists (-*log*_10_ of FDR => 5) **(Fig. 4C)**.

**Figure 4.**
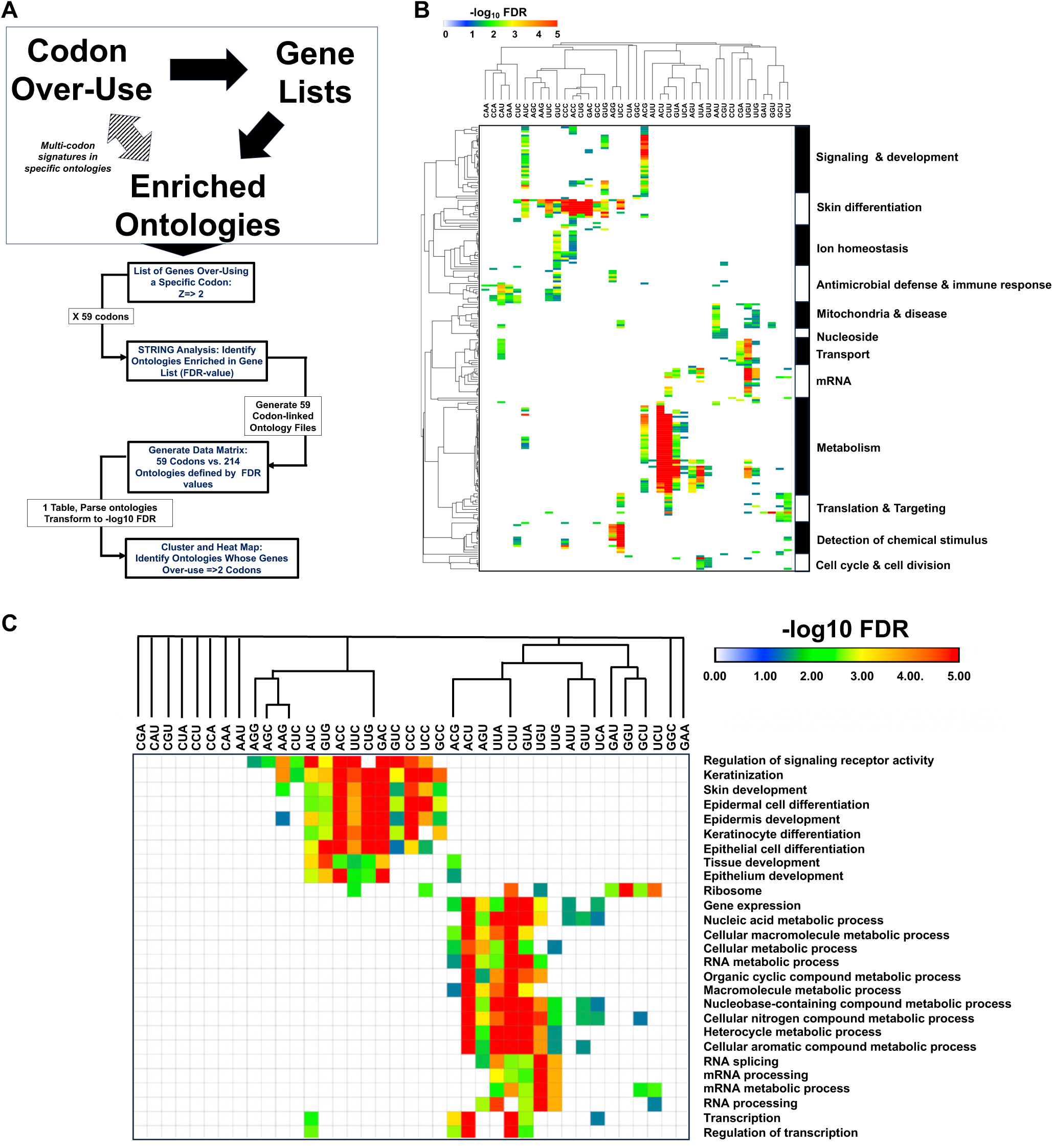
Specific codon biases are linked to distinct biological functions in humans. **(A)** methodology to (1) link genes that over-use a codon with biological process and (2) identify biological process linked multiple codons. **(B)** Gene ontology enriched (FDR < 0.05, -log_10_ FDR-values > 1.3) in each list of codon-biased genes (Z => 2) was identified for 59 codons. Ontologies not found were assigned -log_10_ FDR -values = 0. Data was hierarchically clustered and visualized. Summarized ontologies are listed on the Y-Axis, with exact ontologies shown in supplementary figure S2. **(C)** Data from panel B was filtered to identify ontologies with at least 5 codon linked -log_10_ FDR-values > 2.

**Figure 5.**
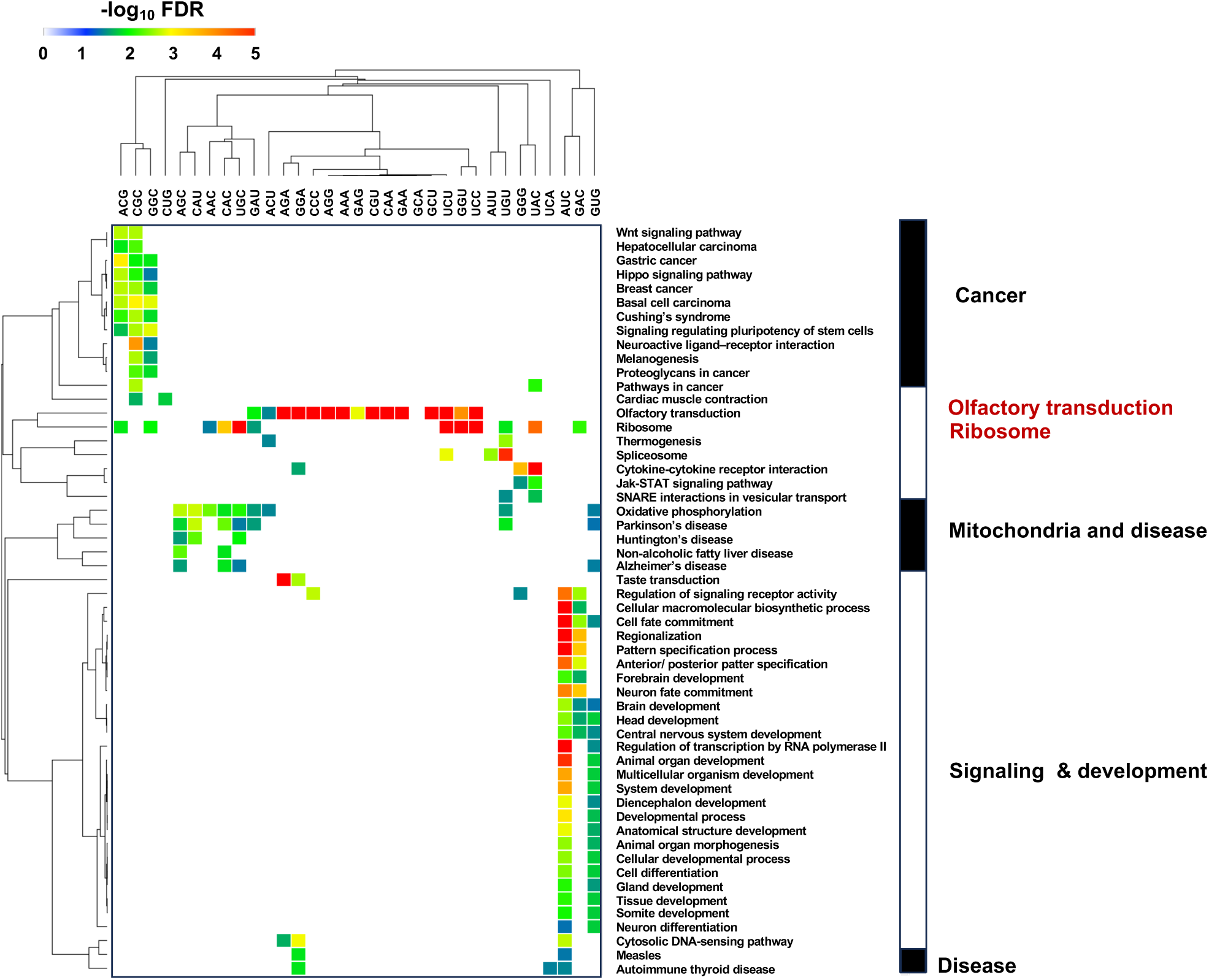
Specific codon biases are linked to distinct biological functions in mice. Gene ontology annotations enriched (FDR < 0.05, -log_10_ FDR > 1.3) in each list of codon-biased genes (Z => 2) was identified for 59 codons. Data was compiled to identify ontologies similarly identified in multiple codons. General ontology categories are shown on far left in black font, with exact ontologies noted in red font.

**Fig. 6.**
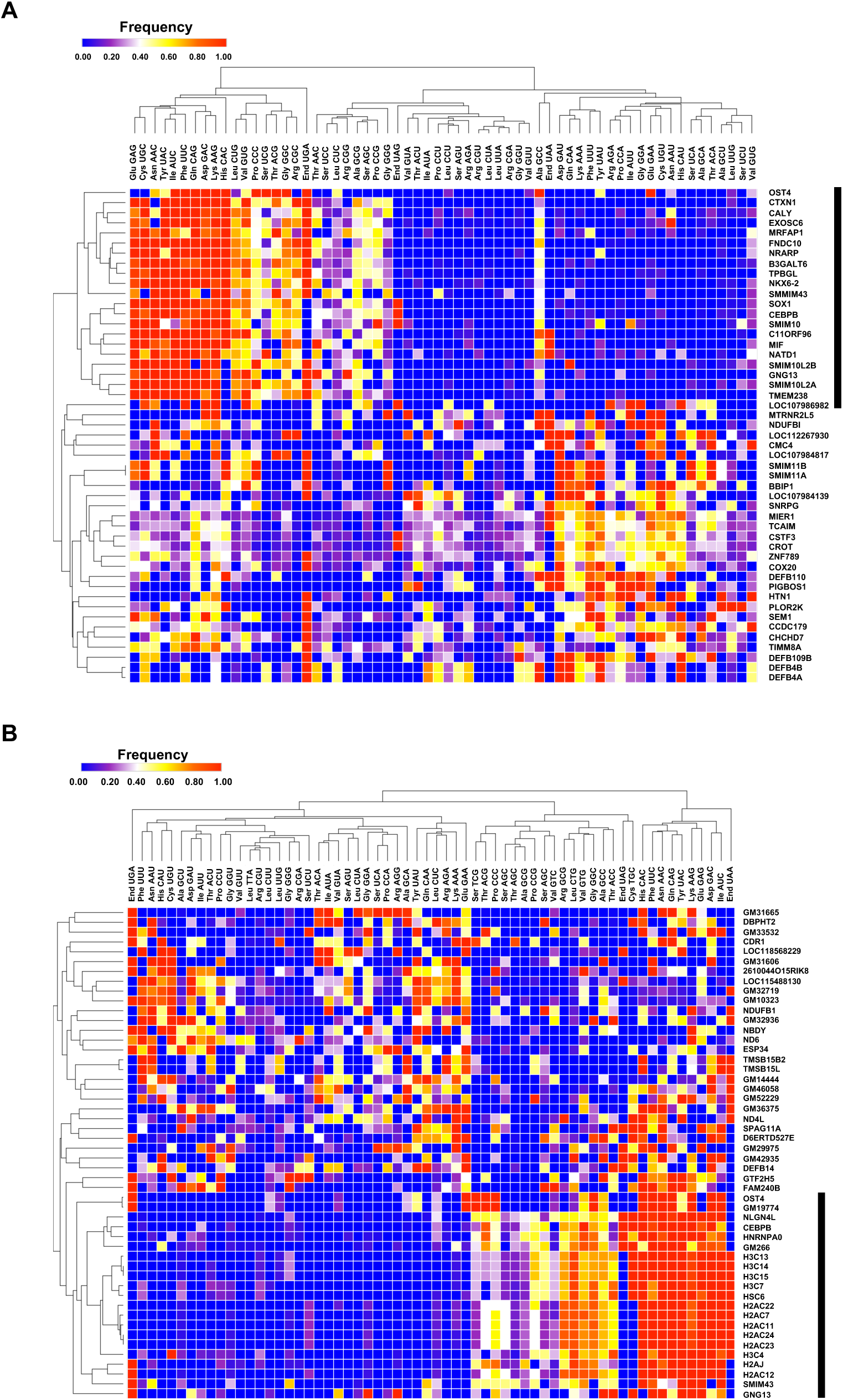
There are two types of extremely biased human and mouse genes. Heat maps detailing the codon frequencies of top 50 biased human (**A**) and mouse (**B**) genes. The black bar denotes clusters populated by genes that over-use synonymous codons mostly ending in G or C.

In mice, ontologies that were identified in at least seven or more lists of genes over-using a specific codon were olfactory transduction, ribosome and oxidative phosphorylation **(Supplemental Table S8B)**. Clusters linked to cancer, olfactory transduction, ribosome, mitochondria & disease, macromolecular processes, signaling & development and diseases were identified **(Fig. 5, Supplemental Table S8B)**. Overall, the number of mouse ontologies having genes that over-use a codon was much less than observed in humans **(Fig. 5 vs Fig. 4)**. For mice, the olfactory transduction ontology is comprised of genes that when combined over-use 15 individual codons, which greatly exceeds the 4 over-used codons found in the identical ontology in humans. Olfactory transduction is represented by genes that over-use 11 codons corresponding to -log_10_ of FDR values => 5, the maximum value on the heat map, and include AGA (Arg), GGA (Gly), CCC (Pro), AGG (Arg), AAA (Lys), CGU (Arg), CAA (Gln), GAA (Glu), GCU (Ala), UCU (Ser), and AUU (Ile). Olfactory transduction is the most codon-distinct ontology identified in mice.

### Some human and mouse genes are extremely biased for many codons

After identifying ontologies that are linked to multiple codons, we investigated if specific genes would have distinct codon bias signatures. ORFs with a high GSZ-score over- and under-use multiple codons and represent the most biased genes in the genome. GSZ-scores and distributions were plotted for human and mouse genes **(Supplemental Table S9A-B, Supplemental Fig. 3A-B).** The top 1% and 2.5% of the codon biased genes were subjected to gene ontology (GO) analysis and the extreme codon biased genes in humans are linked to transcriptional regulation and mitochondria **(Supplemental Table 10A-B)**. In mice the extreme codon biased genes are linked to the spliceosome, beta defensins and defense response to bacterium **(Supplemental Table 10C-D)**. We also analyzed the ICF’s for the top 50 most extreme codon biased human and mouse genes using hierarchical clustering, with the corresponding heat maps identifying two clusters (C/G-ending vs. U/A-ending) in each species (**Fig. 6A-B).** The most striking clusters in each species (**Fig. 6A-B, black bars)** are highlighted by some genes that are completely committed to specific synonymous codons, being that they only use one codon from the synonymous options for some amino acids. We identified 6 genes in humans (*CTXN1*, *FNDC10*, *SMIM10L2A*, *TMEM238*, *SOX1*, *C11ORF96*) that are completely committed to specific synonymous codons [CAG (Gln), GAG (Glu), UAC (Tyr) AUC (Ile), UGC (Cys), UUC (Phe)] for each of 6 amino acids in their gene sequence, with 14 other genes 90% committed to using these 6 codons. In mice, 9 genes encoding histone proteins are 100% committed to a single codon for each of 6 amino acids [UUC (Phe), CAC (His), GAC (Asp), CAG (Gln), AUC (Ile), GAG (Glu)].

### Codon re-engineering can regulate the levels of specific transcription factors

The amino acids Asn, Asp, His and Tyr are each specified by two codons (C- or U-ending) that are decoded by tRNAs that contain a wobble queosine (Q34). *CEPBP* is totally committed to AAC (Asn), GAC (Asp), CAC (His), and UAC (Tyr) codons in both humans and mice (**Fig. 7A)**. CEBPB is a transcription factor that regulates immune and inflammatory responses, with the conserved codon usage between human and mouse genes suggesting a regulatory role. While *CEBPB* is representative of genes over-using codons that end in C or G, the top 50 most biased genes in both organisms also have entries, represented by *MIER1,* that can be generalized as over-using codons that end in U or A. MIER1 is a transcriptional regulator that is a homolog of the mesoderm induction early response protein characterized in *Xenopus laevis*. Mouse and human *MIER1* have similar codon usage patterns, and both are very committed to CUA (6 of 7 His codons), UAU (11 of 15 Tyr codons), GAU (33 of 42 Asp codons) and AAU (18 of 23 Asn codons) **(Fig. 7B)**. We tested the regulatory effects of extreme codon bias in human *CEBPB* and *MIER1*, with a focus on the Asn, Asp, His and Tyr codons. Human *CEBPB* is completely committed to the C-ending codons for Asn (AAC), Asp (GAC), His (CAC) and Tyr (UAC), and contains 41 in total. We generated a synthetic gene for *CEBPB*, (*41Q-CEBPB*) that had the 41 C-ending codons changed to 41 U-ending counterparts, which will produce identical proteins at the amino acid level while testing the importance of codon usage on output. After transfecting the plasmid that expressed *CEBPB* variants, we analyzed for protein levels relative to an internal control **(Fig. 7C)**. We show that under untreated conditions protein levels that are translated from WT *CEBPB* that contains 41 C-ending codons are significantly higher than those translated from the 41Q version that has 41 U-ending codons. A similar trend was observed after NaAsO_2_ treatment, with a slight increase observed for WT and slight decrease observed for 41Q. In contrast to *CEBPB*, *MIER*1 is very committed to U-ending codons for Asn, Asp, His, and Tyr, and contains 68 in total **(Fig. 7D)**. We constructed completely committed C-ending (Q-UP) and U-ending (Q-DW) versions of *MIER1* for Asn, Asp, His and Tyr, and analyzed protein levels. Relative to WT MIER1, the Q-UP version had increased protein levels while Q-DW had decreased protein levels **(Fig. 7D)**. We also performed RNA structure calculations for WT and codon engineered constructs. Minimum Free Energy (MFE) values calculated by RNAfold, UNAfold, and SRNA show similarities between CEPBP-WT and CEPBP-41Q constructs **(Supplementary Figure S4-5)**, with the MIER1-QUP construct having a slightly lower MFE (higher structure) value than MIER-1WT and MIER1-QDW. Together, these results support the idea that C-ending codons in native *CEBPB* and re-engineered *MIER1* promote translation and increased protein levels, with U-ending codons having the opposite effect.

**Figure. 7.**
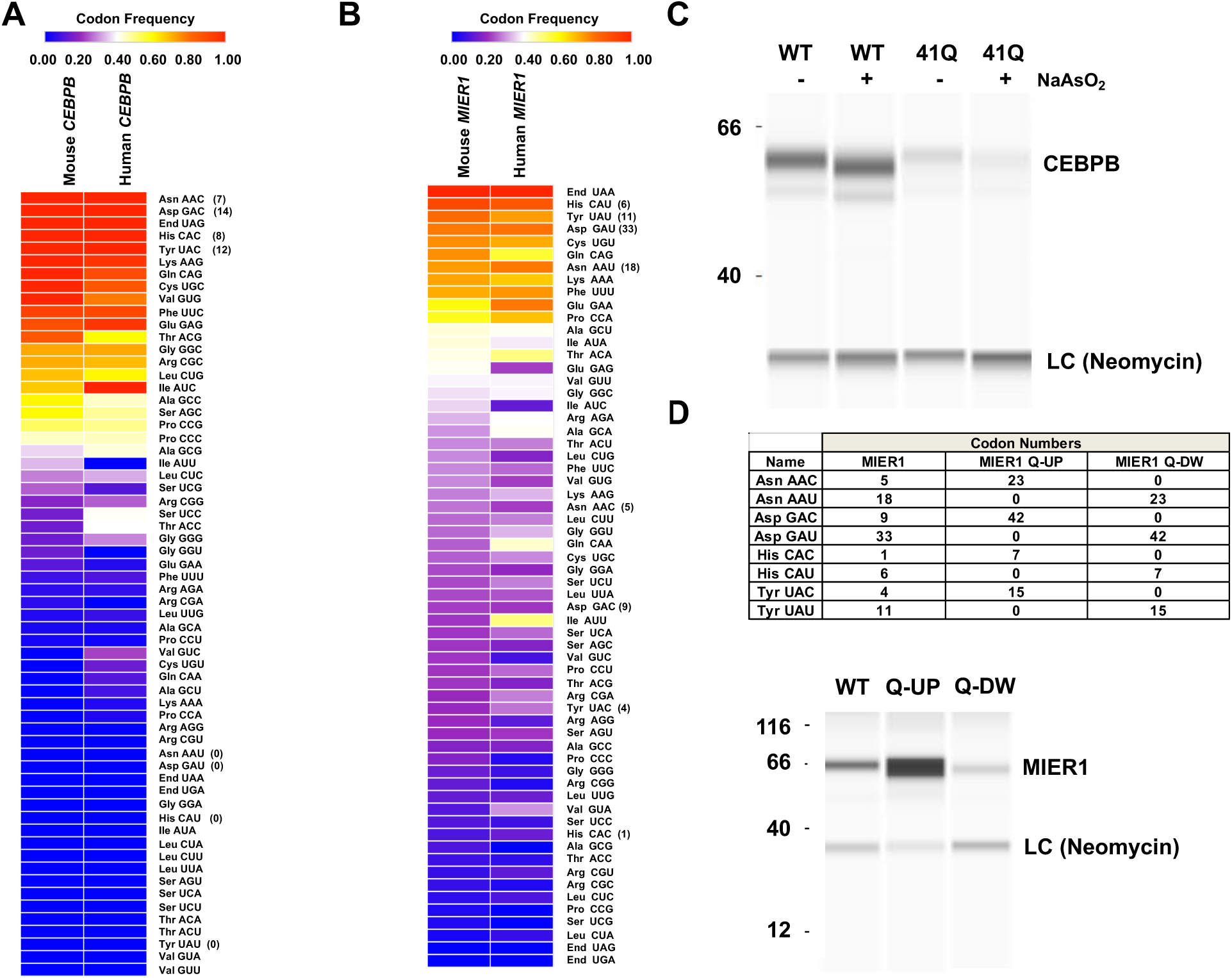
Extreme codon bias can be conserved and regulates protein levels of the CEBPB and MIER1 transcription factors. Heat map-based comparison of gene-specific codon frequencies for human and mouse (A) *CEBPB* and (B) *MIER1* genes. Codons marked with an asterisk are decoded by Q, with parenthesis indicating the number of times that codon is found in the native gene sequence. **(C)** Protein-simple based analysis of WT and re-engineered human CEBPB. **(D)** Codon details (upper) and Protein-simple based analysis (lower) of WT and re-engineered human *MIER1* versions.

## Discussion

### Humans and mice use codon bias in distinct gene families that are potential MoTTs

Previous studies have identified species-specific codon signatures in MoTTs encoding DNA damage response, ROS-detoxification, ribosome and translation-related process and signal transduction pathways.^7,13–15,17,22,23,26,27,37^ ICF values do not report on codon bias, but clustering highlights distinct patterning with underlying over- or under-usage bias. ICF patterns distinctly stratified clusters of mouse genes enriched for biological process of gamete generation, sexual reproduction, spermatid development, defense response to bacterium and innate immune response (all FDR-values < 5E^-21^). Clustering based on ICF values for humans identified ontologies that included chemical stimulus, development (nervous system, system, animal organ), regulation of transcription and neurogenesis (all FDR-values < 5E^-13^). While ICF clustering highlights codon-based differences between mice and humans, development related ontologies were similarly identified in both species, with some distinctions. Species-specific distinctions were most evident using codon over-use ontology mapping, with this methodology utilizing codon bias. Humans genes belonging to ontologies linked to skin differentiation were unique when compared to mice and over-used many codons. Human genes linked to skin differentiation are likely MoTTs. Development has previously been shown to be under extensive translational regulation by Sshu, Wnt, Hippo, PI3K and MAPK pathways.^38^ In addition, upstream – ORF (uORF) mediated translational regulation has been linked to neurogenesis,^38^ and there could be overlapping translation initiation and elongation programs that utilize uORFS and codon usage. Codon usage can also clearly influence mRNA stability, with codon over-use potentially having direct and indirect effects on translation.^39,40^ In mice, the ontology of olfactory transduction is unique compared to humans and it is populated by genes that over-using multiple codons and encode likely MoTTs. Mice have evolved new gene lineages that include those linked to reproduction, immunity, and olfaction,^41^ with the last likely due to rodents’ extensive use of smell. Shared codon usage in pathways has the potential to coordinate the regulation of pathways and also promote complex formations.^42^ Translational regulation of human mRNAs linked to skin development and mouse mRNAs linked to smell could be used to quickly adapt to changing environments, via translational regulation of existing mRNAs, which may be accompanied by dynamic changes in tRNA modifications.

### Different multi-codon over-use signatures are present in human and mouse biological processes

Different types of codon bias have been linked to the regulation of mRNAs whose corresponding proteins belong to specific biological processes in bacteria, yeast, mice and cancer models. In mice, the response to oxidative stress is regulated by increased mcm^5^Um34 modifications that promote the decoding of UGA codons for selenocysteine and found in the mRNAs for ROS-detoxification enzymes.^14^ In humans, METTL1-dependent methylation of G47 to N7-methylguanosine (m^7^G) occurs in multiple tRNAs to increase abundance, with m^7^G on tRNA Arg-TCT-4-1 in cancers promoting the increased translation of mRNAs for cell cycle regulators enriched in the correspondingly decoded AGA codon for Arg.^28,43^ The theme of codon bias regulating stress responses is conserved across phylogeny, with each species utilizing distinct codon – tRNA modification rules. Some specificity in how each species utilizes codon bias should be expected as codon usage in bacteria, yeast and mammals is notably different, which can be attributed to a range of GC contents, from 38% to 67%.^44–46^ In addition, the physiological needs and environmental conditions of each species is different and have likely provided pressure to regulate pathways with different types of codon bias. These species-specific examples demonstrate some simple codon – tRNA modification rules, that are likely more complex in mechanism. METTL1 modifies 25 distinct tRNAs charged with 16 different amino acids and has been linked to stem cell development, cancer, and aging.^28,43^ METTL1’s substrates highlight the potential complexity of identifying codon over-use patterns. Similarly the ELP writers modify tRNA isoacceptors, decoding 13 codons linked to 6 amino acids (Arg, Gly, Glu, Lys, Gln, Sec), with ELP linked to codon biased translational regulation of stress and chemotherapeutic resistance programs in cancers.^8^ We identified complex patterns of codon over-use in our study, with more codons linked to more ontologies in humans then mice. In humans, ontologies linked to skin differentiation and metabolism show two distinct clusters, highlighting multi-codon signatures around groups of codons predominantly ending in C/G or U/A, respectfully. For example, the ontology of keratinization is comprised of genes that over-use 12 C/G ending-codons. In contrast the ontology of cellular nitrogen compound metabolic processing, which was similar to nucleic acid metabolic process, is comprised of groups of genes over-using mostly U- or A-ending codons. Patterning of mRNAs based on codons that end in either U/A or C/G has been identified during responses to stress,^12,47^ and is likely due to specific translational programs that optimize the decoding of corresponding mRNAs. The mouse ontology of olfactory transductions is comprised of mRNAs that as a group over-use 15 specific codons and represent the most complex pattern identified in our study. Olfactory transduction genes over-use 11 codons [AGA (Arg), GGA (Gly), AGG (Arg), AAA (Lys), GAG (Glu), CGU (Arg), CAA (Gln), GAA (Glu), and GGU (Gly)] that are decoded by tRNAs containing the wobble uridine modifications mcm^5^U, mcm^5^s^2^U or mchm^5^U. These wobble uridine modifications in mice are written by Elp1-6, Ctu1-2 and Alkbh8, which would suggest a role for these writers in regulating smell. The mouse olfactory transduction genes also over-use CCC (Pro), GCU (Ala), UCU (Ser), and UCC (Ser) codons, implicating other tRNAs and writers in translational regulation.

### Extreme codon bias is a regulatory feature in mammalian ORFeomes

While complex codon over-usage signatures define specific pathways, extreme codon usage is evident in specific genes. CEBPB is transcriptional regulator of immune and inflammatory networks and promotes drug resistance in non-small cell lung cancer via regulation by Nrf2.^48–51^ The *CEBPG* gene was totally committed to C-ending codons for Asn, Asp, His and Tyr in humans and mice. Codon re-engineering of *CEBPB* highlights the importance of C-ending codons in maintaining protein levels. Total conservation between humans and mice supports a regulatory role for extreme codon usage, with corresponding Asn, Asp, His, and Tyr codons all decoded by tRNAs containing the modification Q34. Formation of the Q modification in tRNA of humans is dependent on the diet and microbiome-based production of the essential co-factor queuine,^52,53^ with nutrient levels also affecting Q-levels in Trypanasomes.^54^ Regulation of immune and inflammatory networks in both humans and mice may be tied to diet and the microbiome, via tRNA modification and codon-dependent translational regulation. The human transcription factors NKX6-2 and SOX1 are also highly committed to C-ending codons decoded by Q (NKX6-2 has 33 and SOX1 has 54) and use few U-ending codons (NKX6-2 uses 0 and SOX1 uses 1) and these proteins can control cell migration and invasion in gastric cancer cells,^55^ and oncogenic activity in glioblastomas,^56^ respectively. There could thus be therapeutic potential to target the translation of these extreme codon patterns to treat cancers. *MIER1* is a mirror image of *CEBPB*, *NKX6-2,* and *SOX1* mRNA, as it is over-using U-ending codons for Asn, Asp, His and Tyr codons. Codon re-engineering of *MIER1* supports that U-ending codons lead to lower protein levels in the context of the human HEPG2 system. The presence of many genes that over-use codons that end in U (and A) is evident in humans and mice. Translational programs that favor decoding of U-ending codons could be present in certain cell types or physiological conditions, as there is evidence of translational regulation during development and tissue and age-specific gene-expression has been identified.^57–59^ Also, most mitochondrial genes over-use codons that end in U or A, which could sync the translation of U- or A-ending codons in cytoplasmic mRNAs with mitochondrial physiology.

Our studies support an emerging theme that codon bias can be used to regulate transcription factors. Previously codon reengineering of the master transcriptional regulator of dormancy, DosR, in bacteria has been demonstrated to enhance it translation during hypoxia, with the increased wobble U tRNA modification 5-oxyacetyl-uridine (cmo^5^U) postulated to promote ACG decoding, while restricting ACC.^13^ In BRAFV600E-expressing human melanoma cells the translation of AAA (Lys), CAA (Gln) and GAA (Glu) codons in *HIF1A* mRNA has been linked to the wobble U writers for corresponding tRNAs.^8^ HIF1A is part of the transcriptional regulator HIF1 that is induced in response to hypoxia and allows cells to adapt to low oxygen conditions.^60,61^ The transcription factor DEK1 has also been shown to be regulated by the codon biased translation of the LEF1, to coordinate a pro-invasion program in cancer.^62,63^ Codon re-engineering of *HIF1A* and *LEF1* has also be shown to alter their regulation. Codon-bias in the ORFs of transcription factors or their regulatory proteins could allow for translational regulation to control the transcription of regulons.

Altogether, our study has provided evidence of extensive codon bias in specific biological processes for humans and mice. Codon-biased translational regulation has emerged as an important regulatory mechanism in stress responses and human cancers and aging, with strong connections to tRNA writers.^28,43^ Developing approaches to regulate the translation of codon-biased MoTTs could be used to regulate transcription, treat cancers or improve aging, with our study highlighting protein networks with distinct codon patterns and writers to targets. Our findings also provide a blue print to improve the production of proteins with extremely biased Q-decoded codons, with potential applications to improve protein production for biomanufacturing.

## Acknowledgments

The authors are grateful for support from the National Institutes of Health (ES026856, ES031529, GM070641, CA274603) and the National Research Foundation of Singapore through the Singapore-MIT Alliance for Research and Technology Antimicrobial Resistance Interdisciplinary Research Group, the MIT - Spain "la Caixa" Foundation Seed Fund, and the Agilent Foundation. The graphical abstract was generated using BioRender.com.

## Supplemental Tables

**Supplemental Table S1.** Species Specific Codon Data. **(A)** Codon count, ICF, and Z-score data for 19,711 human genes. **(B)** Codon count, ICF, and Z-score data for 22,138 mouse genes. *<u>(These files are too large and will be deposited online)</u>*.

**Supplemental Tables S2.** Gene Lists for Ontology Analysis. Human **(A-B)** and mouse **(C-D)** genes over-represented and specific to Supplemental Table 3. Human **(E-F)** and mouse **(G-H)** genes under-represented and specific to Supplemental Table 3. *<u>(These files are too large</u> and will be deposited online)*.

**Supplemental Table S3A.** Biological processes enriched in 398 human genes over-using up to 12 codons (Z=> 2).

| **Process** | **FDR** |
| --- | --- |
| Mitochondrial respiratory chain complex assembly | 0.00047 |
| Mitochondrion organization | 0.00057 |
| Mitochondrion | 7.85e-05 |
| Cellular protein-containing complex assembly | 0.0139 |

**Supplemental Table S3B.**
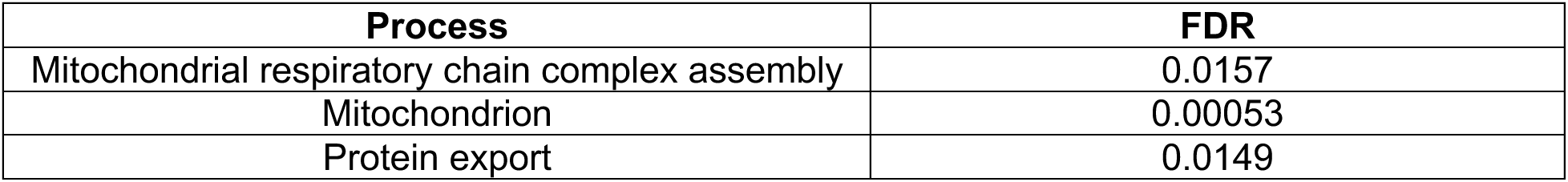
Biological processes enriched in 202 human genes under-using up to 10 codons (Z=> 2).

**Supplemental Table S3C.** Biological processes enriched in 367 mouse genes over-using up to 12 codons (Z=> 2).

| **Process** | **FDR** |
| --- | --- |
| Mitochondrial respiratory chain complex assembly | 0.0442 |
| Mitochondrial protein complex | 0.0171 |
| Parkinson's disease | 0.0090 |
| Beta defensins | 1.36e-05 |

**Supplemental Table S3D.** Biological processes enriched in 367 mouse genes under-using up to 11 codons (Z<= - 2).

| **Process** | **FDR** |
| --- | --- |
| Mitochondrial respiratory chain complex assembly | **0.0046** |
| Defense response to bacterium | 0.0018 |
| Ribosome | 0.0021 |
| Parkinson's disease | 0.0357 |

**Supplemental Table S4.** Data for Figure 1. **(A)** Human data for figure 1D. **(B)** Human data for figure 1E. **(C)** Mouse data for figure 1F. **(D)** Mouse data for figure 1G. *<u>(These files are too large and will be deposited online)</u>*.

**Supplemental Tables S5.** ICF Data used for clustering. **(A)** Human and **(B)** mouse ICF data. *(These files are too large and will be deposited online)*.

**Supplemental Tables S6.**
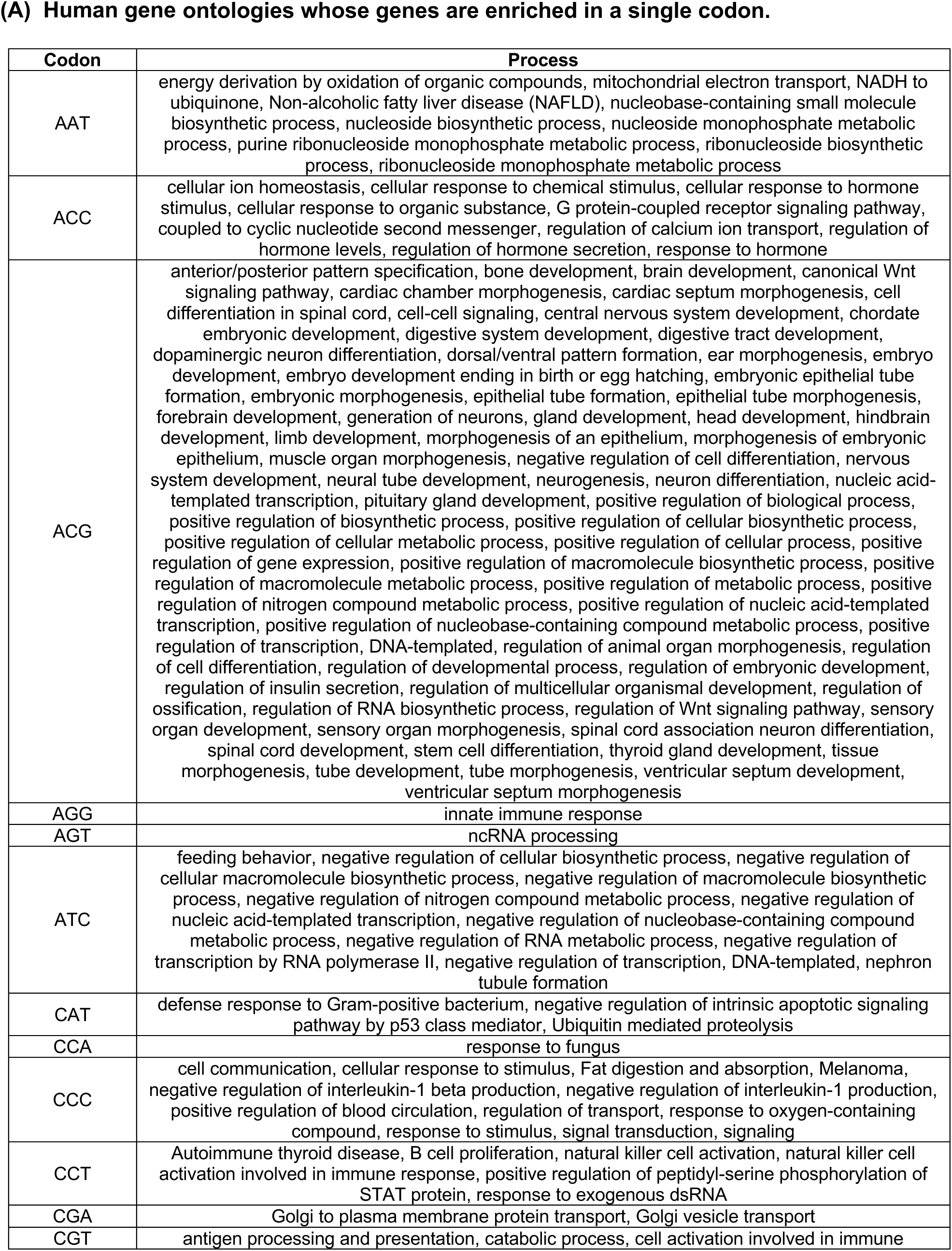
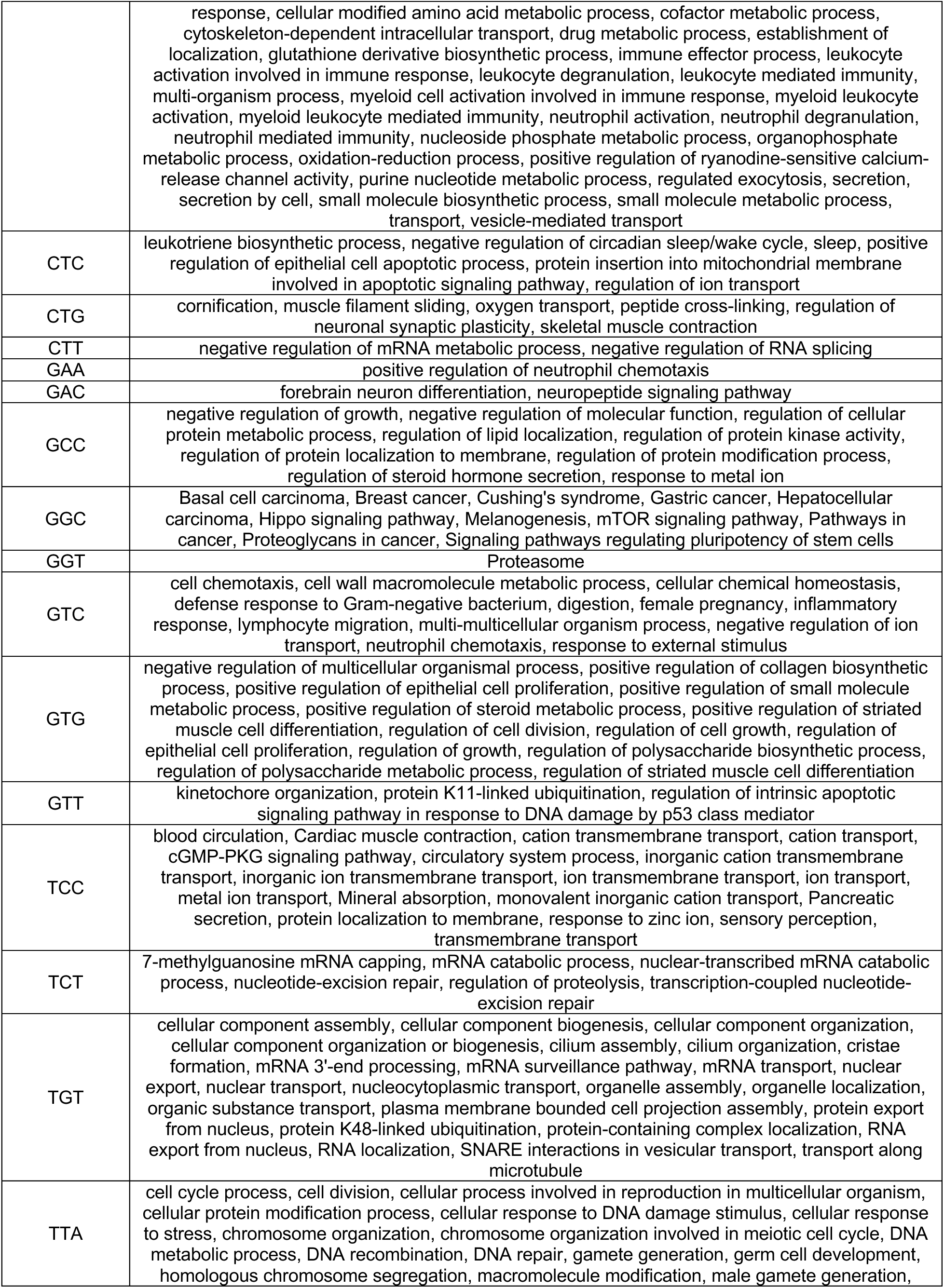
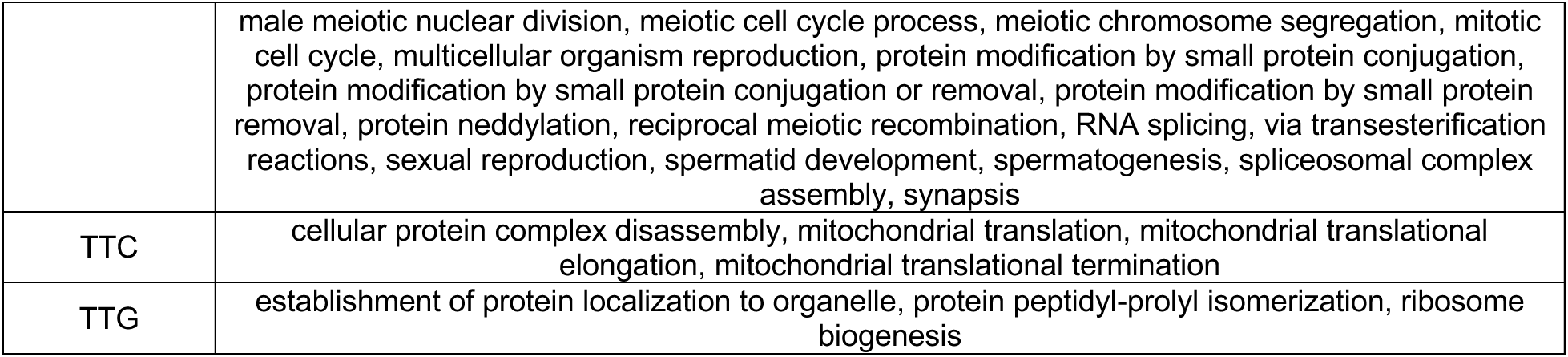
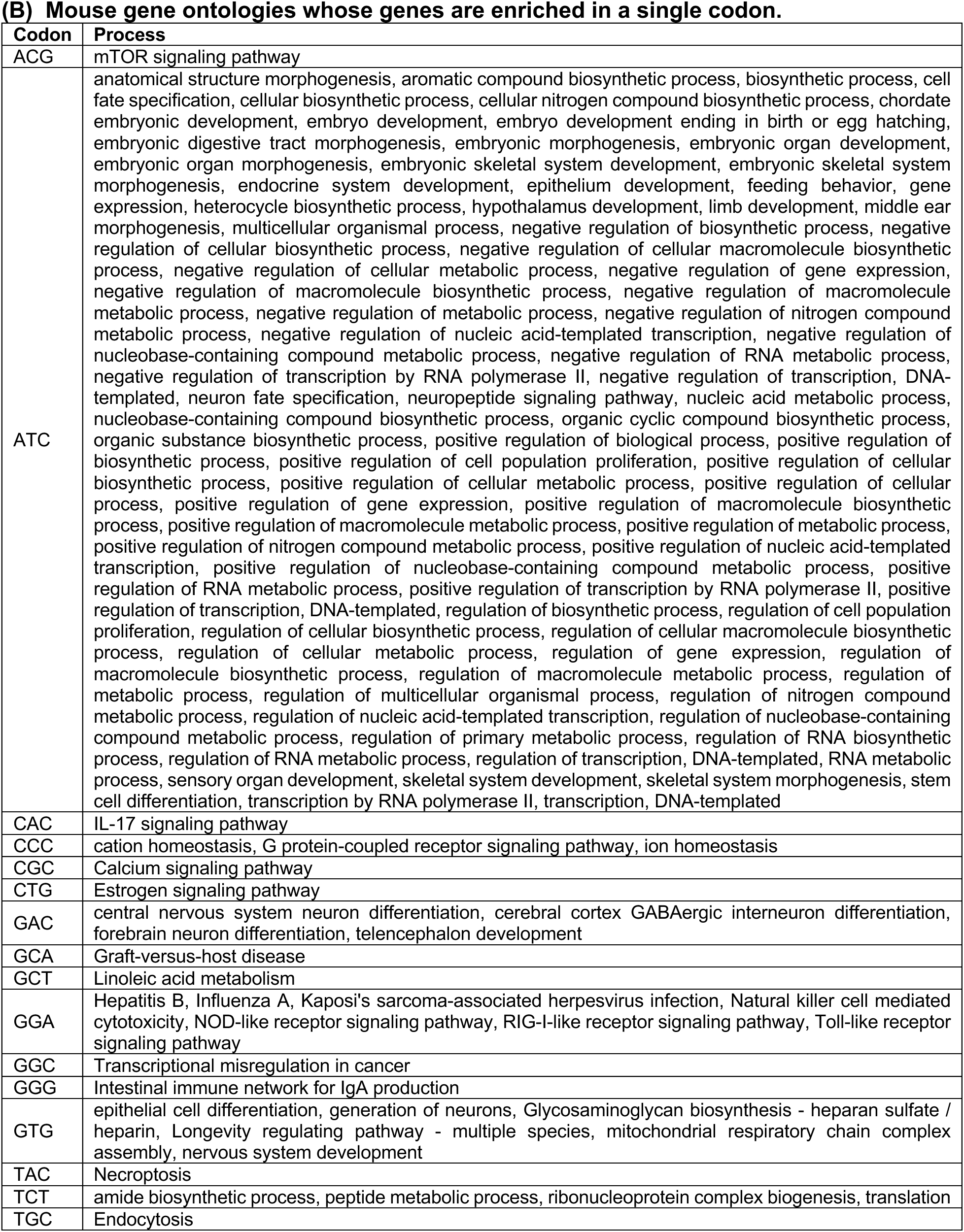
Ontologies enriched in a single ontology.

| Codon | Process |
| --- | --- |
| AAT | energy derivation by oxidation of organic compounds, mitochondrial electron transport, NADH to ubiquinone, Non-alcoholic fatty liver disease (NAFLD), nucleobase-containing small molecule biosynthetic process, nucleoside biosynthetic process, nucleoside monophosphate metabolic process, purine ribonucleoside monophosphate metabolic process, ribonucleoside biosynthetic process, ribonucleoside monophosphate metabolic process |
| ACC | cellular ion homeostasis, cellular response to chemical stimulus, cellular response to hormone stimulus, cellular response to organic substance, G protein-coupled receptor signaling pathway, coupled to cyclic nucleotide second messenger, regulation of calcium ion transport, regulation of hormone levels, regulation of hormone secretion, response to hormone |
| ACG | anterior/posterior pattern specification, bone development, brain development, canonical Wnt signaling pathway, cardiac chamber morphogenesis, cardiac septum morphogenesis, cell differentiation in spinal cord, cell-cell signaling, central nervous system development, chordate embryonic development, digestive system development, digestive tract development, dopaminergic neuron differentiation, dorsal/ventral pattern formation, ear morphogenesis, embryo development, embryo development ending in birth or egg hatching, embryonic epithelial tube formation, embryonic morphogenesis, epithelial tube formation, epithelial tube morphogenesis, forebrain development, generation of neurons, gland development, head development, hindbrain development, limb development, morphogenesis of an epithelium, morphogenesis of embryonic epithelium, muscle organ morphogenesis, negative regulation of cell differentiation, nervous system development, neural tube development, neurogenesis, neuron differentiation, nucleic acid-templated transcription, pituitary gland development, positive regulation of biological process, positive regulation of biosynthetic process, positive regulation of cellular biosynthetic process, positive regulation of cellular metabolic process, positive regulation of cellular process, positive regulation of gene expression, positive regulation of macromolecule biosynthetic process, positive regulation of macromolecule metabolic process, positive regulation of metabolic process, positive regulation of nitrogen compound metabolic process, positive regulation of nucleic acid-templated transcription, positive regulation of nucleobase-containing compound metabolic process, positive regulation of transcription, DNA-templated, regulation of animal organ morphogenesis, regulation of cell differentiation, regulation of developmental process, regulation of embryonic development, regulation of insulin secretion, regulation of multicellular organismal development, regulation of ossification, regulation of RNA biosynthetic process, regulation of Wnt signaling pathway, sensory organ development, sensory organ morphogenesis, spinal cord association neuron differentiation, spinal cord development, stem cell differentiation, thyroid gland development, tissue morphogenesis, tube development, tube morphogenesis, ventricular septum development, ventricular septum morphogenesis |
| AGG | innate immune response |
| AGT | ncRNA processing |
| ATC | feeding behavior, negative regulation of cellular biosynthetic process, negative regulation of cellular macromolecule biosynthetic process, negative regulation of macromolecule biosynthetic process, negative regulation of nitrogen compound metabolic process, negative regulation of nucleic acid-templated transcription, negative regulation of nucleobase-containing compound metabolic process, negative regulation of RNA metabolic process, negative regulation of transcription by RNA polymerase II, negative regulation of transcription, DNA-templated, nephron tubule formation |
| CAT | defense response to Gram-positive bacterium, negative regulation of intrinsic apoptotic signaling pathway by p53 class mediator, Ubiquitin mediated proteolysis |
| CCA | response to fungus |
| CCC | cell communication, cellular response to stimulus, Fat digestion and absorption, Melanoma, negative regulation of interleukin-1 beta production, negative regulation of interleukin-1 production, positive regulation of blood circulation, regulation of transport, response to oxygen-containing compound, response to stimulus, signal transduction, signaling |
| CCT | Autoimmune thyroid disease, B cell proliferation, natural killer cell activation, natural killer cell activation involved in immune response, positive regulation of peptidyl-serine phosphorylation of STAT protein, response to exogenous dsRNA |
| CGA | Golgi to plasma membrane protein transport, Golgi vesicle transport |
| CGT | antigen processing and presentation, catabolic process, cell activation involved in immune |
|  | response, cellular modified amino acid metabolic process, cofactor metabolic process, cytoskeleton-dependent intracellular transport, drug metabolic process, establishment of localization, glutathione derivative biosynthetic process, immune effector process, leukocyte activation involved in immune response, leukocyte degranulation, leukocyte mediated immunity, multi-organism process, myeloid cell activation involved in immune response, myeloid leukocyte activation, myeloid leukocyte mediated immunity, neutrophil activation, neutrophil degranulation, neutrophil mediated immunity, nucleoside phosphate metabolic process, organophosphate metabolic process, oxidation-reduction process, positive regulation of ryanodine-sensitive calcium-release channel activity, purine nucleotide metabolic process, regulated exocytosis, secretion, secretion by cell, small molecule biosynthetic process, small molecule metabolic process, transport, vesicle-mediated transport |
| CTC | leukotriene biosynthetic process, negative regulation of circadian sleep/wake cycle, sleep, positive regulation of epithelial cell apoptotic process, protein insertion into mitochondrial membrane involved in apoptotic signaling pathway, regulation of ion transport |
| CTG | cornification, muscle filament sliding, oxygen transport, peptide cross-linking, regulation of neuronal synaptic plasticity, skeletal muscle contraction |
| CTT | negative regulation of mRNA metabolic process, negative regulation of RNA splicing |
| GAA | positive regulation of neutrophil chemotaxis |
| GAC | forebrain neuron differentiation, neuropeptide signaling pathway |
| GCC | negative regulation of growth, negative regulation of molecular function, regulation of cellular protein metabolic process, regulation of lipid localization, regulation of protein kinase activity, regulation of protein localization to membrane, regulation of protein modification process, regulation of steroid hormone secretion, response to metal ion |
| GGC | Basal cell carcinoma, Breast cancer, Cushing's syndrome, Gastric cancer, Hepatocellular carcinoma, Hippo signaling pathway, Melanogenesis, mTOR signaling pathway, Pathways in cancer, Proteoglycans in cancer, Signaling pathways regulating pluripotency of stem cells |
| GGT | Proteasome |
| GTC | cell chemotaxis, cell wall macromolecule metabolic process, cellular chemical homeostasis, defense response to Gram-negative bacterium, digestion, female pregnancy, inflammatory response, lymphocyte migration, multi-multicellular organism process, negative regulation of ion transport, neutrophil chemotaxis, response to external stimulus |
| GTG | negative regulation of multicellular organismal process, positive regulation of collagen biosynthetic process, positive regulation of epithelial cell proliferation, positive regulation of small molecule metabolic process, positive regulation of steroid metabolic process, positive regulation of striated muscle cell differentiation, regulation of cell division, regulation of cell growth, regulation of epithelial cell proliferation, regulation of growth, regulation of polysaccharide biosynthetic process, regulation of polysaccharide metabolic process, regulation of striated muscle cell differentiation |
| GTT | kinetochore organization, protein K11-linked ubiquitination, regulation of intrinsic apoptotic signaling pathway in response to DNA damage by p53 class mediator |
| TCC | blood circulation, Cardiac muscle contraction, cation transmembrane transport, cation transport, cGMP-PKG signaling pathway, circulatory system process, inorganic cation transmembrane transport, inorganic ion transmembrane transport, ion transmembrane transport, ion transport, metal ion transport, Mineral absorption, monovalent inorganic cation transport, Pancreatic secretion, protein localization to membrane, response to zinc ion, sensory perception, transmembrane transport |
| TCT | 7-methylguanosine mRNA capping, mRNA catabolic process, nuclear-transcribed mRNA catabolic process, nucleotide-excision repair, regulation of proteolysis, transcription-coupled nucleotide-excision repair |
| TGT | cellular component assembly, cellular component biogenesis, cellular component organization, cellular component organization or biogenesis, cilium assembly, cilium organization, cristae formation, mRNA 3'-end processing, mRNA surveillance pathway, mRNA transport, nuclear export, nuclear transport, nucleocytoplasmic transport, organelle assembly, organelle localization, organic substance transport, plasma membrane bounded cell projection assembly, protein export from nucleus, protein K48-linked ubiquitination, protein-containing complex localization, RNA export from nucleus, RNA localization, SNARE interactions in vesicular transport, transport along microtubule |
| TTA | cell cycle process, cell division, cellular process involved in reproduction in multicellular organism, cellular protein modification process, cellular response to DNA damage stimulus, cellular response to stress, chromosome organization, chromosome organization involved in meiotic cell cycle, DNA metabolic process, DNA recombination, DNA repair, gamete generation, germ cell development, homologous chromosome segregation, macromolecule modification, male gamete generation, |
|  | male meiotic nuclear division, meiotic cell cycle process, meiotic chromosome segregation, mitotic cell cycle, multicellular organism reproduction, protein modification by small protein conjugation, protein modification by small protein conjugation or removal, protein modification by small protein removal, protein neddylation, reciprocal meiotic recombination, RNA splicing, via transesterification reactions, sexual reproduction, spermatid development, spermatogenesis, spliceosomal complex assembly, synapsis |
| TTC | cellular protein complex disassembly, mitochondrial translation, mitochondrial translational elongation, mitochondrial translational termination |
| TTG | establishment of protein localization to organelle, protein peptidyl-prolyl isomerization, ribosome biogenesis |
| ACG | mTOR signaling pathway |
| ATC | anatomical structure morphogenesis, aromatic compound biosynthetic process, biosynthetic process, cell fate specification, cellular biosynthetic process, cellular nitrogen compound biosynthetic process, chordate embryonic development, embryo development, embryo development ending in birth or egg hatching, embryonic digestive tract morphogenesis, embryonic morphogenesis, embryonic organ development, embryonic organ morphogenesis, embryonic skeletal system development, embryonic skeletal system morphogenesis, endocrine system development, epithelium development, feeding behavior, gene expression, heterocycle biosynthetic process, hypothalamus development, limb development, middle ear morphogenesis, multicellular organismal process, negative regulation of biosynthetic process, negative regulation of cellular biosynthetic process, negative regulation of cellular macromolecule biosynthetic process, negative regulation of cellular metabolic process, negative regulation of gene expression, negative regulation of macromolecule biosynthetic process, negative regulation of macromolecule metabolic process, negative regulation of metabolic process, negative regulation of nitrogen compound metabolic process, negative regulation of nucleic acid-templated transcription, negative regulation of nucleobase-containing compound metabolic process, negative regulation of RNA metabolic process, negative regulation of transcription by RNA polymerase II, negative regulation of transcription, DNA-templated, neuron fate specification, neuropeptide signaling pathway, nucleic acid metabolic process, nucleobase-containing compound biosynthetic process, organic cyclic compound biosynthetic process, organic substance biosynthetic process, positive regulation of biological process, positive regulation of biosynthetic process, positive regulation of cell population proliferation, positive regulation of cellular biosynthetic process, positive regulation of cellular metabolic process, positive regulation of cellular process, positive regulation of gene expression, positive regulation of macromolecule biosynthetic process, positive regulation of macromolecule metabolic process, positive regulation of metabolic process, positive regulation of nitrogen compound metabolic process, positive regulation of nucleic acid-templated transcription, positive regulation of nucleobase-containing compound metabolic process, positive regulation of RNA metabolic process, positive regulation of transcription by RNA polymerase II, positive regulation of transcription, DNA-templated, regulation of biosynthetic process, regulation of cell population proliferation, regulation of cellular biosynthetic process, regulation of cellular macromolecule biosynthetic process, regulation of cellular metabolic process, regulation of gene expression, regulation of macromolecule biosynthetic process, regulation of macromolecule metabolic process, regulation of metabolic process, regulation of multicellular organismal process, regulation of nitrogen compound metabolic process, regulation of nucleic acid-templated transcription, regulation of nucleobase-containing compound metabolic process, regulation of primary metabolic process, regulation of RNA biosynthetic process, regulation of RNA metabolic process, regulation of transcription, DNA-templated, RNA metabolic process, sensory organ development, skeletal system development, skeletal system morphogenesis, stem cell differentiation, transcription by RNA polymerase II, transcription, DNA-templated |
| CAC | IL-17 signaling pathway |
| CCC | cation homeostasis, G protein-coupled receptor signaling pathway, ion homeostasis |
| CGC | Calcium signaling pathway |
| CTG | Estrogen signaling pathway |
| GAC | central nervous system neuron differentiation, cerebral cortex GABAergic interneuron differentiation, forebrain neuron differentiation, telencephalon development |
| GCA | Graft-versus-host disease |
| GCT | Linoleic acid metabolism |
| GGA | Hepatitis B, Influenza A, Kaposi's sarcoma-associated herpesvirus infection, Natural killer cell mediated cytotoxicity, NOD-like receptor signaling pathway, RIG-I-like receptor signaling pathway, Toll-like receptor signaling pathway |
| GGC | Transcriptional misregulation in cancer |
| GGG | Intestinal immune network for IgA production |
| GTG | epithelial cell differentiation, generation of neurons, Glycosaminoglycan biosynthesis - heparan sulfate / heparin, Longevity regulating pathway - multiple species, mitochondrial respiratory chain complex assembly, nervous system development |
| TAC | Necroptosis |
| TCT | amide biosynthetic process, peptide metabolic process, ribonucleoprotein complex biogenesis, translation |
| TGC | Endocytosis |

**Supplemental Table S7.** Data for Figures 6 and 7. **(A)** Human data for figure 6: GO Map. **(B)** Mouse data for figure 7: GO map. *(These files are too large and will be deposited online)*.

**Supplemental Table S8.**
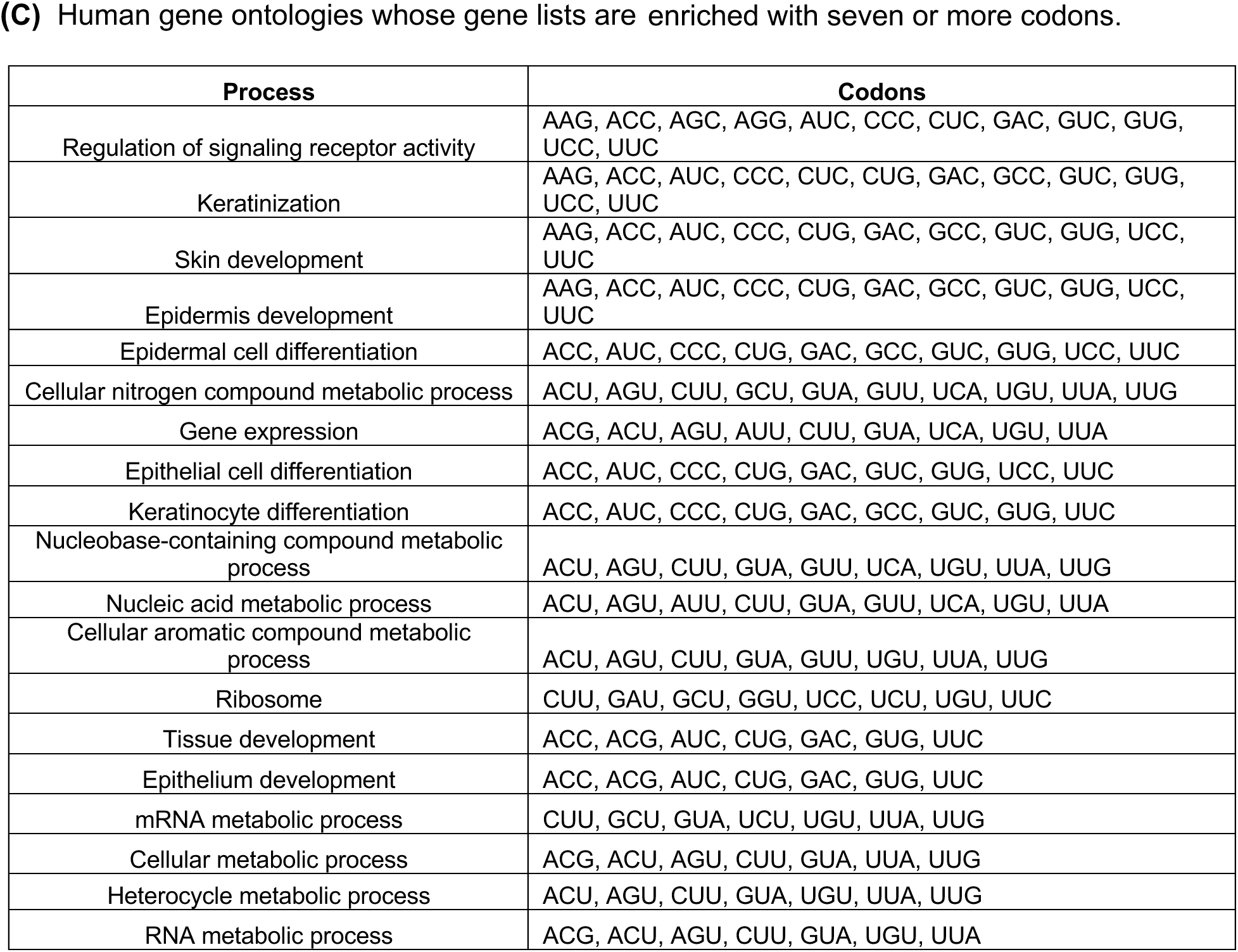
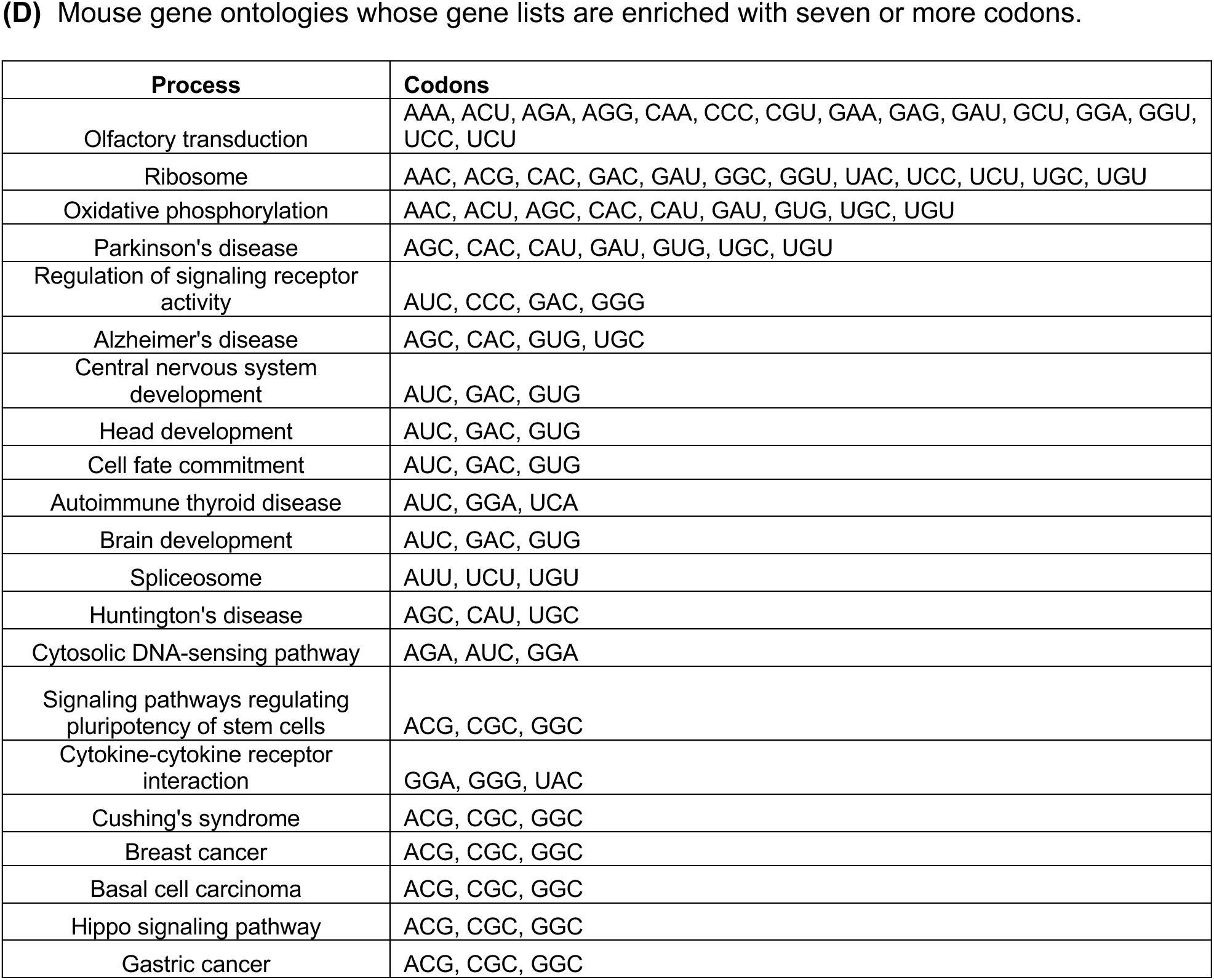
Data files derived from figure 6 and 7. **(A)** All processes with human genes enriched for multiple codons. Human data for figure: GO Map. **(B)** All processes with mouse genes enriched for multiple codons. Mouse data for figure: GO Map. *<u>(These files</u> <u>are too large and will be deposited online).</u>*

| Process | Codons |
| --- | --- |
| Regulation of signaling receptor activity | AAG, ACC, AGC, AGG, AUC, CCC, CUC, GAC, GUC, GUG, UCC, UUC |
| Keratinization | AAG, ACC, AUC, CCC, CUC, CUG, GAC, GCC, GUC, GUG, UCC, UUC |
| Skin development | AAG, ACC, AUC, CCC, CUG, GAC, GCC, GUC, GUG, UCC, UUC |
| Epidermis development | AAG, ACC, AUC, CCC, CUG, GAC, GCC, GUC, GUG, UCC, UUC |
| Epidermal cell differentiation | ACC, AUC, CCC, CUG, GAC, GCC, GUC, GUG, UCC, UUC |
| Cellular nitrogen compound metabolic process | ACU, AGU, CUU, GCU, GUA, GUU, UCA, UGU, UUA, UUG |
| Gene expression | ACG, ACU, AGU, AUU, CUU, GUA, UCA, UGU, UUA |
| Epithelial cell differentiation | ACC, AUC, CCC, CUG, GAC, GUC, GUG, UCC, UUC |
| Keratinocyte differentiation | ACC, AUC, CCC, CUG, GAC, GCC, GUC, GUG, UUC |
| Nucleobase-containing compound metabolic process | ACU, AGU, CUU, GUA, GUU, UCA, UGU, UUA, UUG |
| Nucleic acid metabolic process | ACU, AGU, AUU, CUU, GUA, GUU, UCA, UGU, UUA |
| Cellular aromatic compound metabolic process | ACU, AGU, CUU, GUA, GUU, UGU, UUA, UUG |
| Ribosome | CUU, GAU, GCU, GGU, UCC, UCU, UGU, UUC |
| Tissue development | ACC, ACG, AUC, CUG, GAC, GUG, UUC |
| Epithelium development | ACC, ACG, AUC, CUG, GAC, GUG, UUC |
| mRNA metabolic process | CUU, GCU, GUA, UCU, UGU, UUA, UUG |
| Cellular metabolic process | ACG, ACU, AGU, CUU, GUA, UUA, UUG |
| Heterocycle metabolic process | ACU, AGU, CUU, GUA, UGU, UUA, UUG |
| RNA metabolic process | ACG, ACU, AGU, CUU, GUA, UGU, UUA |
| Olfactory transduction | AAA, ACU, AGA, AGG, CAA, CCC, CGU, GAA, GAG, GAU, GCU, GGA, GGU, UCC, UCU |
| Ribosome | AAC, ACG, CAC, GAC, GAU, GGC, GGU, UAC, UCC, UCU, UGC, UGU |
| Oxidative phosphorylation | AAC, ACU, AGC, CAC, CAU, GAU, GUG, UGC, UGU |
| Parkinson's disease | AGC, CAC, CAU, GAU, GUG, UGC, UGU |
| Regulation of signaling receptor activity | AUC, CCC, GAC, GGG |
| Alzheimer's disease | AGC, CAC, GUG, UGC |
| Central nervous system development | AUC, GAC, GUG |
| Head development | AUC, GAC, GUG |
| Cell fate commitment | AUC, GAC, GUG |
| Autoimmune thyroid disease | AUC, GGA, UCA |
| Brain development | AUC, GAC, GUG |
| Spliceosome | AUU, UCU, UGU |
| Huntington's disease | AGC, CAU, UGC |
| Cytosolic DNA-sensing pathway | AGA, AUC, GGA |
| Signaling pathways regulating pluripotency of stem cells | ACG, CGC, GGC |
| Cytokine-cytokine receptor interaction | GGA, GGG, UAC |
| Cushing's syndrome | ACG, CGC, GGC |
| Breast cancer | ACG, CGC, GGC |
| Basal cell carcinoma | ACG, CGC, GGC |
| Hippo signaling pathway | ACG, CGC, GGC |
| Gastric cancer | ACG, CGC, GGC |

**Supplemental Table S9.** Identification of extremely codon biased genes. **(A)** 19,711 summed codon biased human gene scores. **(B)** 22,138 summed codon biased mouse gene Scores. *<u>(These files are too large and will be deposited online)</u>*.

**Supplemental Table S10.**
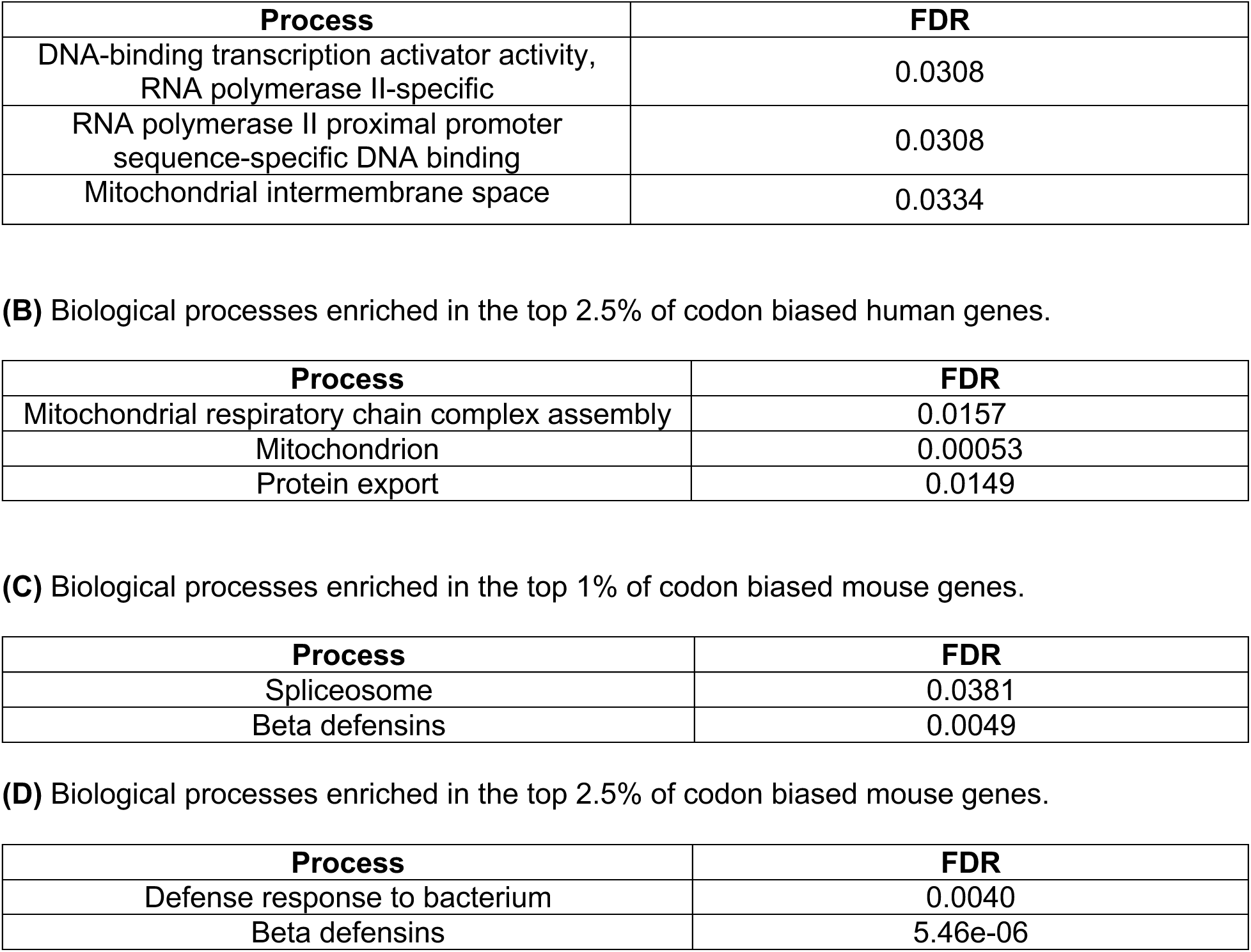
Ontologies of extremely codon biased genes. (A) Top 1% of codon biased human genes.Supplemental Figures

## Supplemental Figures

**Supplemental Figure S1.**
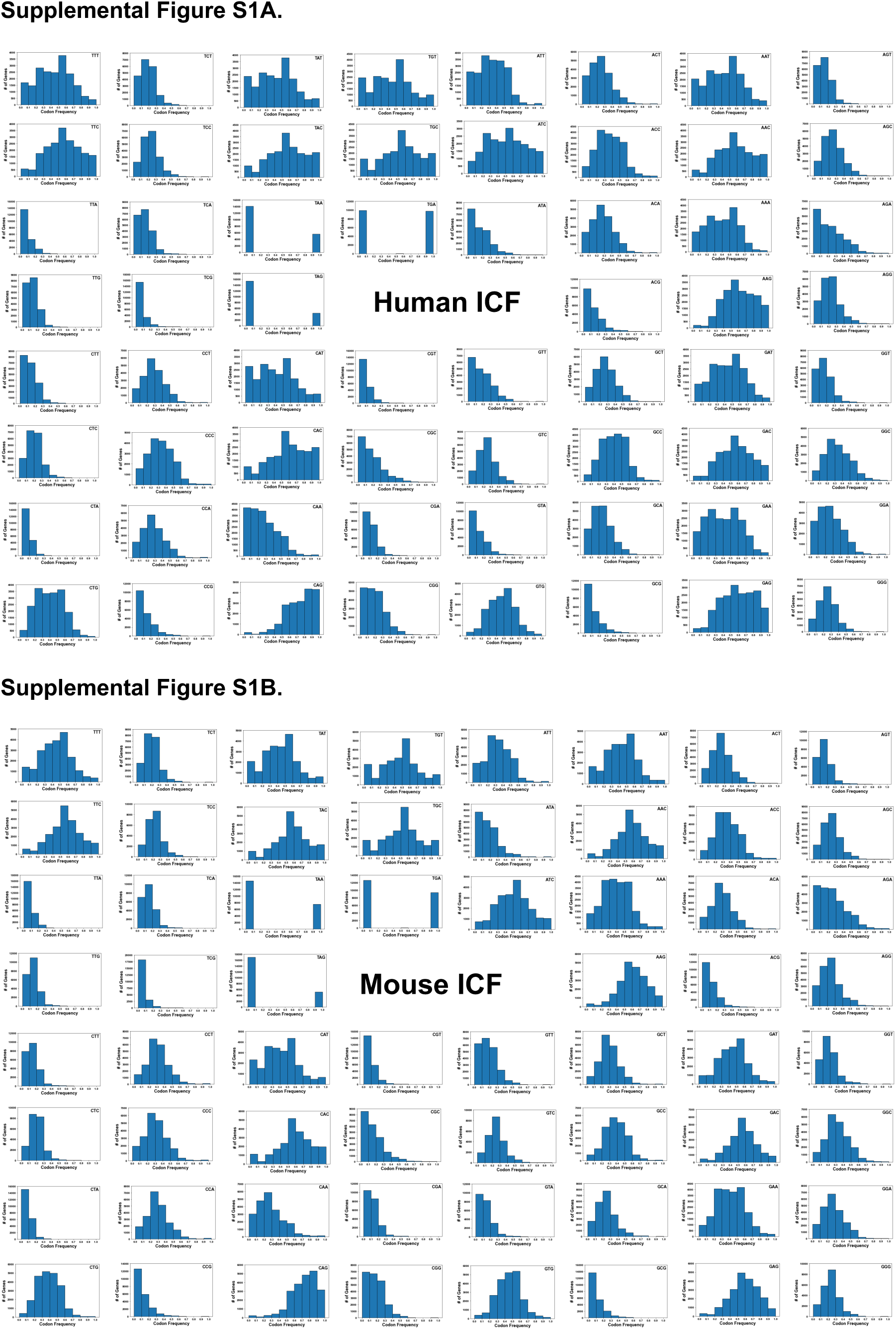
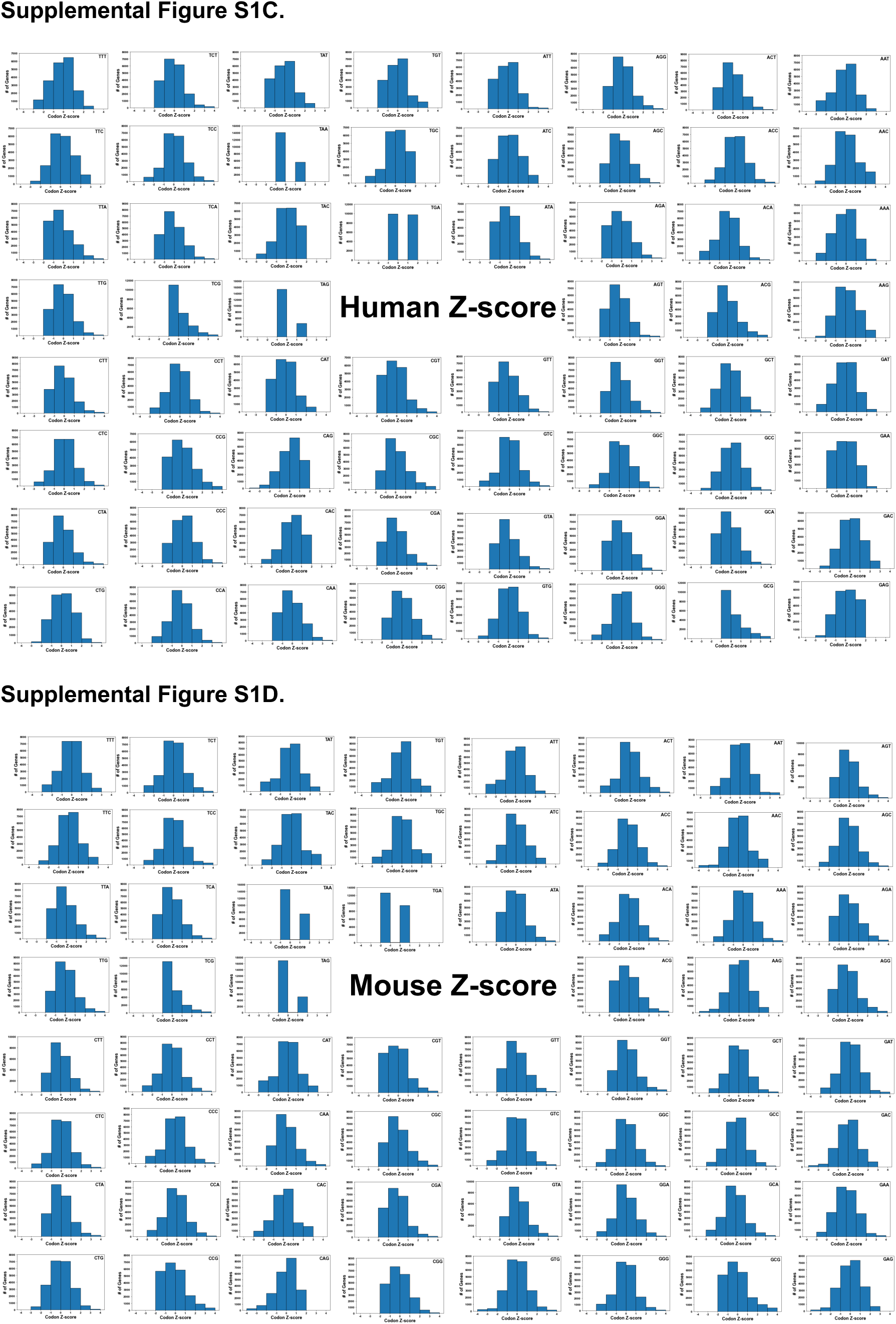
Gene Specific Codon Data in Humans and Mice. (A) Codon ICF histograms for human codon usage. **(B)** Codon ICF histograms for all mouse codon usage. (**C)** Z-score histograms for human codon usage**. (D)** Z-score histograms for mouse codon usage.

**Supplemental Figure S2.**
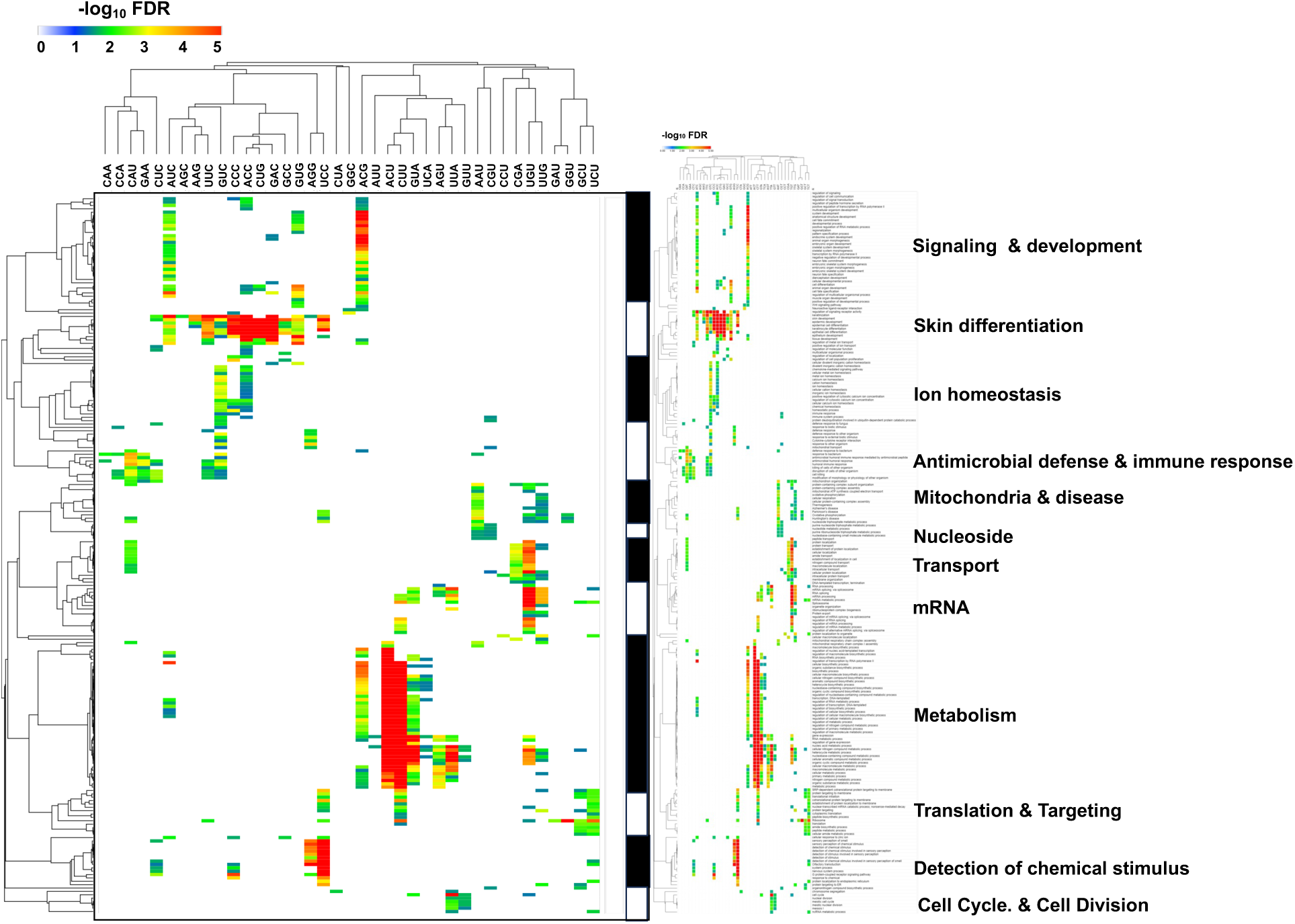
Human Heat Map from figure 4, with all ontologies. Gene ontologies enriched (FDR < 0.05, -log_10_ FDR-values > 1.3) in each list of codon-biased genes (Z => 2) was identified for 59 codons. Ontologies not found were assigned -log_10_ FDR -values = 0. Data was hierarchically clustered and visualized using a heat map to identify ontologies linked to multiple codons (=> 2). Summarized ontologies are listed on the Y-Axis, with exact ontologies also on Y-axis.

**Supplemental Figure S3.**
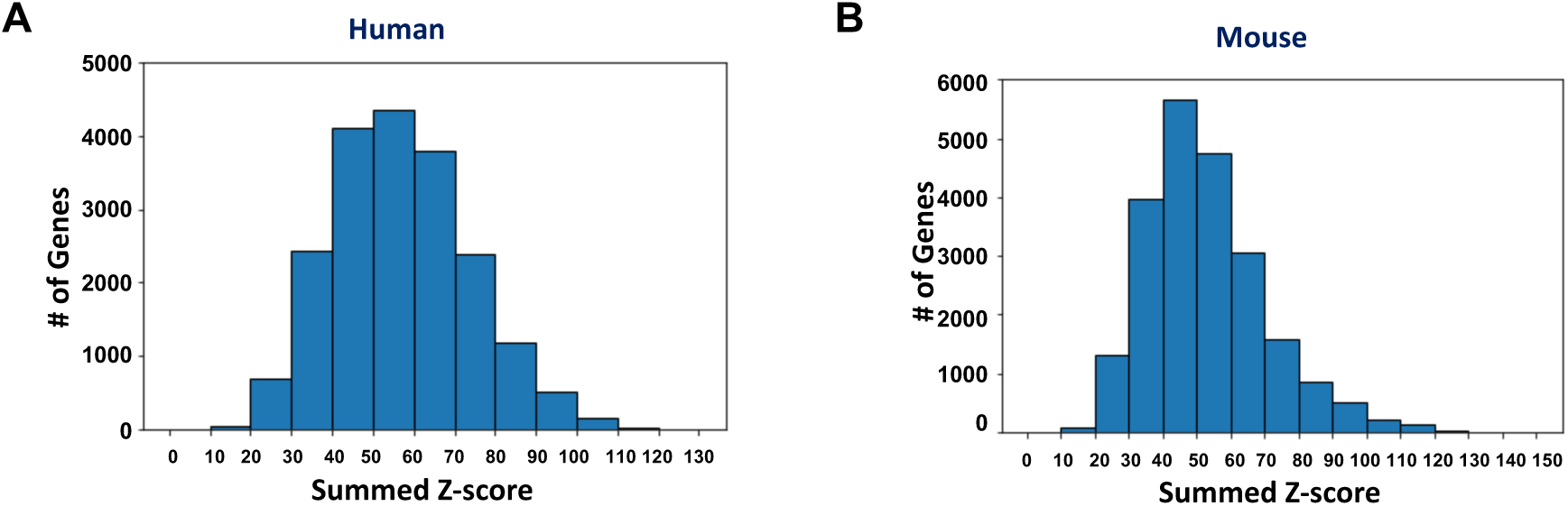
Gene Summed Z-score Values. **(A)** Human and **(B)** mouse GSZ-score distributions.

**Supplemental Figure S4.**
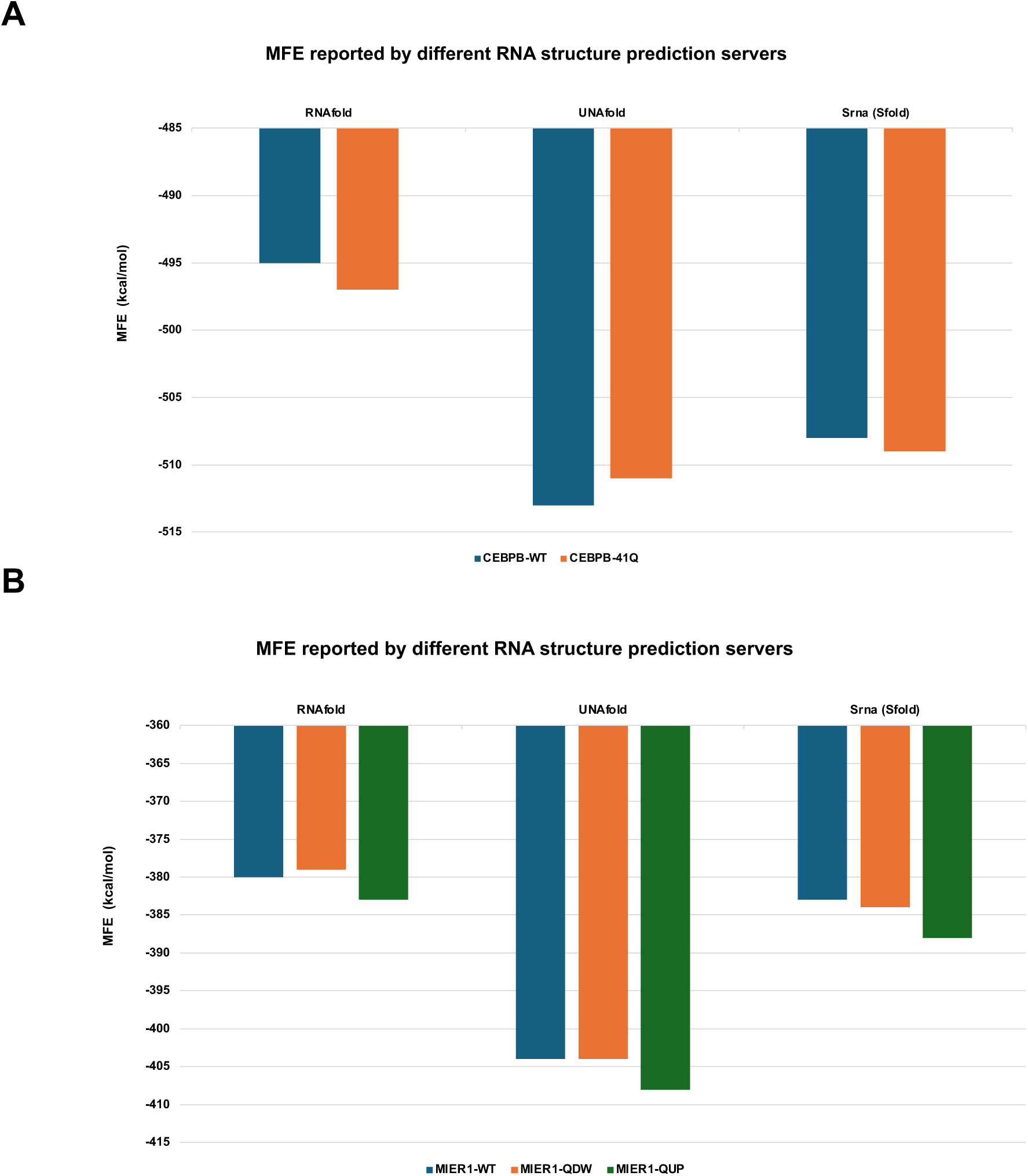
mRNA Stability Measures. **(A)** MFE comparisons between RNAfold, UNAfold, and SRNA for WT and engineered sequences. The plot of the MFE values predicted by the three different tools show similarities between CEPBP-WT and CEPBP-41Q constructs and **(B)** MIER-WT, MIER1-QUP and MIER1-QDW.

**Supplemental Figure S5.**
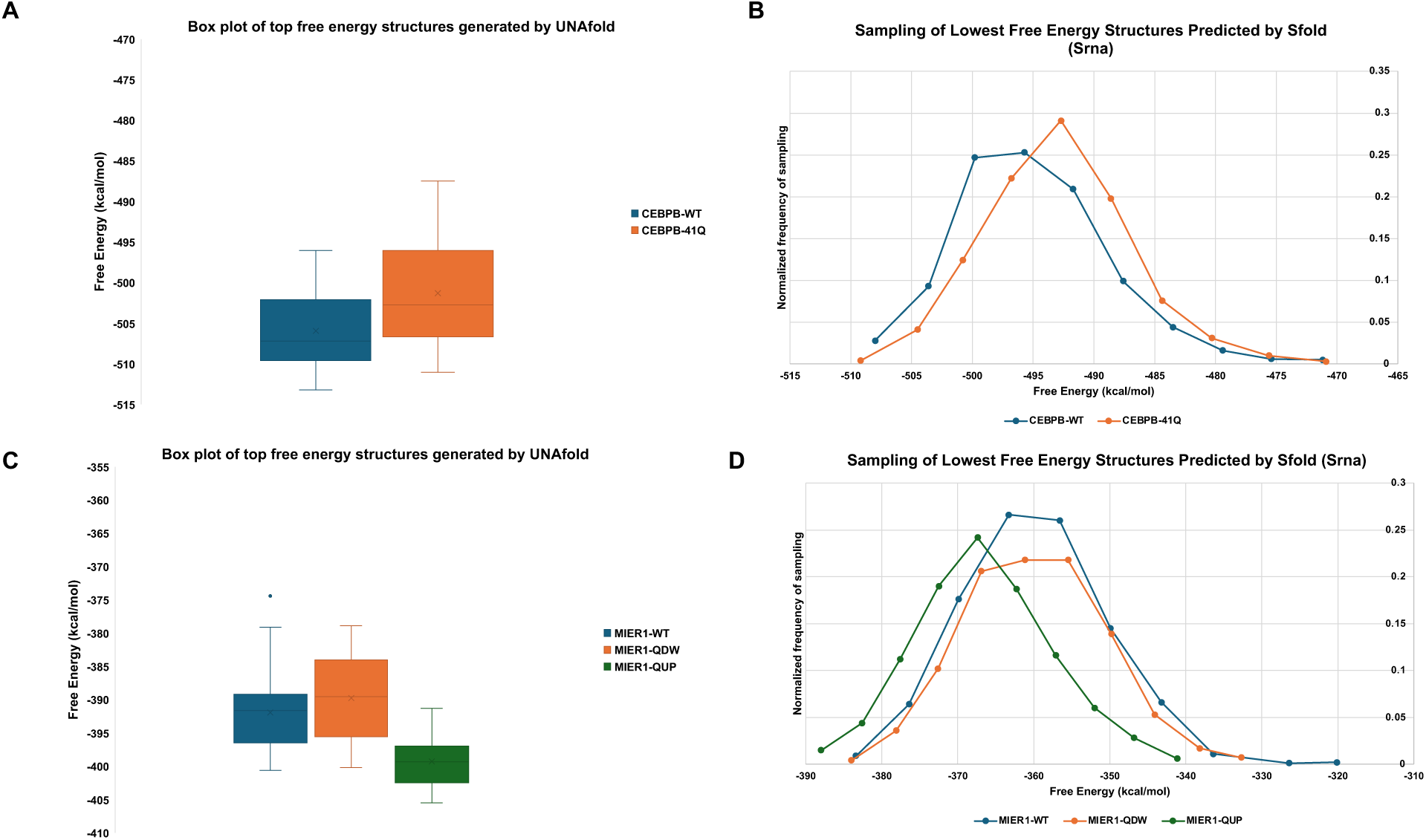
RNA structure calculations for WT and codon engineered constructs. When the energies of top structures with lowest free energy obtained from (A) UNAfold and **(B)** SFold were compare. CEPBP-WT displayed a lower free energy (higher structure) than CEPBP-41. **(C)** The free energy structures sampled by MIER1-QDW were increased when compared to MIER-WT and **(D)** those sampled by MIER1-QUP showed a lower free energy trend when compared to MIER1-WT and MIER1-QDW. In general, the prevalence of higher structure element in RNA and lower free energies for CEPBP-WT and MIER1-QUP correlates with observed higher protein output.

## Notes

### Competing Interest Statement

The authors have declared no competing interest.

